# Neural markers for Musical Creativity in Jazz Improvisation and Classical Interpretation

**DOI:** 10.1101/2021.02.12.427648

**Authors:** Shama Sarwat Rahman, Kim Christensen, Henrik Jeldtoft Jensen, Peter Vuust, Joydeep Bhattacharya

**Affiliations:** Imperial College London, Centre for Complexity Science, South Kensington Campus, London SW7 2AZ; Aarhus University, Centre for Music in the Brain, Nørrebrogade 44 building NBG/ 10G, 10G-5-29, 8000 Aarhus C, Denmark; Goldsmiths University, Centre for Cognition, Culture and Computation, New Cross, London SE14 6NW

**Keywords:** Creativity, Improvisation, Interpretation, EEG, sLORETA, Musical

## Abstract

Two main types of musical creativity in the western canon are improvisation and interpretation. With improvisation, the fundamental structure of the melody, chords, rhythm and tempo of a piece can be modified, while with interpretation, the focus is on the emotional dynamics. Here we characterise electrical brain activity from professional jazz and classical pianists, whilst they were engaged in these different creative tasks with musical excerpts from both genres. Multivariate EEG was recorded during two phases of each task, mental planning and actual performance. Subsequently neuronal activity patterns were source localised with standardised low resolution electromagnetic tomography (sLORETA). For each musical performance, we obtained both subjective (self-rated) and objective (blind, expert-rated) measures of musical creativity. Across both tasks and genre backgrounds, within the first and middle 4 second segments of the performance phase, for musical performances that were judged highly creative objectively by external expert music assessors, we observed an increased activation in the anterior cingulate and medial prefrontal cortex (Brodmann area, BA32), suggesting a maintenance of executive control, and integrating motoric and emotional communication during creativity. Across genre backgrounds, within the performance phase for the interpretation task compared to the improvisation task, there was an increased activity in the insula (BA 13), suggesting a convergent creative task from the linked goal-orientated conscious error-monitoring and audio-visual integration functions. Genre profession also gave rise to differences across phases; jazz pianists presented a decreased parietal (BA7) activity during improvisation tasks suggesting a role for defocussed attention and for classical pianists, both tasks were associated with occipitotemporal (BA 37) activity which is involved in semantic/ metaphorical processing suggesting a close adherence to the visual score. These 3 areas relate the cognitive demands of the creative musical task to the demands of the corresponding genre of music.

**Highlights:**

- EEG activity associated to musical creativity types: Improvisation and Interpretation
- Increased activity in Insula (BA 13) for Interpretation suggest convergent creativity
- Decreased Precuneus (BA7) activity for Improvisation suggest defocussed attention
- Increased activity in medial prefrontal cortex (BA32) in highly creative performance

## INTRODUCTION

What is creativity? What mental processes and cognitive pathways are involved? Are there distinct neurobiological correlates?

By an unique tripartite approach of scaled self-assessments and objective expert ratings which are matched to EEG recordings, our research attempts to characterise creativity by identifying neural patterns and correlates in brain activity which typically appear during the most creative performances. The focus of the study is particularly on music.

We wished to investigate the following hypothesis that there are differences in brain activity that are indicative of creativity, and moreover the type of creativity undertaken whilst being reflective of the genre of music. Moreover, the temporal evolution present in both musical improvisation and interpretation would also be represented in brain activity. We also expected there could typically be a misfit between subjective self-assessments and independent expert judges’ assessments and thus objective EEG measures would most likely correlate with judges’ assessments as compared to and over self-assessments.

Traditionally, experimental studies investigating creativity properties have focused on problem-solving using mental exercises, mind games and brain teasers derived for the purposes of a psychological study e.g. Guildford’s Alternative Uses Task (1967) or Torrance Tests of Creative Thinking (1962). Additionally, qualitative models of creativity have been proposed, some defining the stages of problem-solving such as Wallas’ model or mental models such as the Geneplore Model which divides creativity into a generative and exploratory phase [Finke et al 1992]. We wish to extend the body of work in the field of creativity by exploring a real-world situation such as the creative performance of music using the apt techniques of improvisation or interpretation. Like talent, creativity is dependent on domain-specific knowledge [Weisberg, 1999]. As such, the focus of this research is specifically on musical creativity for the purposes of a controlled experimental paradigm that can shed light on general creativity across different disciplines, as they may share a certain basic process.

We wished to explore the extent to which the psychological creative process correlates with neurophysiologically measurable signatures. To contrast different types of creative music making we compared Classical and Jazz musicians and their creative performances of excerpts from these different musical genres whilst engaged in both improvisation and interpretation.

*Improvisation* refers to the ability of the performer to change the structure of a musical phrase by modifying its key, melodic contour, the notes, rhythm and time signature. The improviser may seem to have an unlimited set of choices but they are not necessarily unconstrained. Musical improvisation does implicitly depend on a specific musical style, and therefore, is constrained by the rules and constraints of that musical style e.g. orthodox modern jazz [Johnson-Laird, 2002]. Jazz offers a great model for improvisational creativity as the written outline of the music is often sketchy and its artistic success relies on the performers ability to improvise over a combination of the melody and a given chord scheme [Berliner, 1994; Vuust, 2009; Monson, 1997].

Improvisation involves a wide range of complex cognitive processes along with strong emotional components as “the improvisers must effect real-time sensory and perceptual coding, optimal attention allocation, event-interpretation, decision making, prediction, memory storage and recall, error correction, and movement control, and further, must integrate these processes into an optimally seamless set of musical statements that reflect both a personal perspective on musical organisation and a capacity to affect listeners” [Pressing, 1998]. In fact, improvisation can be likened to real-time composition where a musical phrase is generated from the mind perhaps from a theme or accompanied by visual imagery which is a form of mental models which simulate processes in the real-world.

*Interpretation* refers to the ability of the performer to interpret the composer’s markings of dynamics, tempo and emotionality without changing the written score in their performance. It involves being creative within the given structure of the musical phrase without modifying its key, melodic contour, the notes, rhythm and time signature. This skill is found most commonly in western classical music where musicians have a strong focus on shaping the finer aspects of musical phrasing and personal interpretation. As such, classical music is a good model for interpretational creativity and by comparing these 2 genres we can look into the deeper intricacies of musical creativity.

Furthermore, studying a performing musician in action offers an unique opportunity to investigate human creativity as it unfolds in real-time. Due to the temporal evolution in improvisation [Sawyer, 1992] and interpretation [Dean and Balles, 2010], and also the particular global structure of the music the performer may create or phrase [Cooper and Meyer, 1960], this may also be reflected in large-scale brain activity. As EEG has a high temporal resolution, it is advantageous over other techniques such as fMRI and PET studies in melody and sentence generation [Brown, Martinez and Parsons, 2006], to observe and characterise how the different regions of the brain interact during the rapid creative process.

A few studies have investigated the brain responses of creative professionals in action. During mental composition of drawings, professional artists showed higher long-range synchronization in the delta band (1–4 Hz), thereby indicating enhanced top-down processing in artists; non-artists, on the other hand, showed more local synchronization indicative of bottom-up processing [Bhattacharya and Petsche, 2005]. Emphasized top-down processing is also observed in professional dancers during mental imagery of an improvised dance but not during a learned dance routine [Fink *et al*., 2009].

Both of these studies though not specific to music, use EEG. An fMRI study [Limb and Braun, 2008] reported that during musical improvisational accompaniment, a widespread brain network gets activated but concurrently the dorsolateral prefrontal cortex, a crucial brain region involved with planning and self-monitoring, gets deactivated. As the study used accompaniment as a basis for improvisation, the nature of the task is more representative of interpretative goal-oriented creativity. Furthermore, the significant role of memory cannot be ruled out.

In another fMRI study, Bengtsson and colleagues [Bengtsson et al, 2007] investigated piano improvisation, by employing three experimental conditions of *improvise, reproduce* and *free improvisation*. A broad network of brain regions, including sensorimotor cortex (presupplementary motor area, the rostral part of the dorsal premotor cortex), superior temporal gyrus, and the prefrontal cortex, specifically the right DLPFC were found to be associated with piano improvisation. However, the all-male participants were presented with musical excerpts that were in one of only two keys (Fminor or Fmajor), and asked to lay supine in the fMRI machine (as they were in the Limb and Braun study) to play on a small keyboard with only their right hand. The authors also query whether the difference in the ‘Improvise*-*Reproduce’ comparison could be down to a non-accurate replication during Reproduce resulting in different motor outputs not attributable to the cognitive demands of improvisation; this replication again introduces the additional role of memory.

Drawing from and adding to the conclusions of this prior research, using EEG as a technique would lead to better ecological validity. Inside an fMRI scanner, the pianists were asked to play whilst lying down, which might have involved different motor skills and cause different perceptions and reactions than usual, as pianists normally perform sitting upright. EEG does not pose such limitations and though head movements have to be kept to a minimum, the technique allows us to get closer to observing this creative activity without affecting it in the process.

Furthermore, on the subject of judging creativity, both Braun *et al*. (2012) who investigated verbal creativity with lyrical rapping, and Villareal *et al*. (2013) who investigated rhythmical musical creativity, made use of a compendium of creativity assessments posed only to judges and not to the musicians themselves. The former, rather than directly addressing the fundamental essence of creativity, employed a dossier of questions that gathered evidence of creativity from associated musical properties (e.g., “Did the participant vary rhythmic patterns?”), and the latter employing a marketing technique called SCAMPER that measures the effect of creativity (an acronym for “substitute, combine, adapt, modify, put to other uses, eliminate and reverse”). Judging the extent to which a certain performance can be considered creative is not straight-forward and there are no universally accepted creativity tests with the majority of functional neuroimaging studies exploring creativity using different methods of assessment [Treffinger, 1986] and focussing more on the personality traits associated with the creativity ‘*syndrome*’ [Runco, 2004]. In our experimental design, we circumvented the reliance on creativity tests through an inclusion of musicians’ opinions themselves on what constitutes a creative performance in addition to collecting and matching these with assessments of experts in the field (building on Amabile’s Consensual Assessment Technique (1982)).

Our study has the dimension of physical performance (as opposed to only mental imagery), as we reason that this actualization is vital to the coherent cognitive processing of music to result in and during a creative state for a performer. In addition, our study focuses on improvisation that is more free and compositional in nature, rather than as accompaniment, and does not have the added cognitive complication of memory in EEG signatures as participants are shown musical excerpts (as stimuli) that they are unfamiliar with using handpicked music manuscripts of Classical music from the 15th century to present from the British Library and commissioning new Jazz compositions, standardising both into unique scores readable for musicians from both backgrounds. Importantly, we have tried staying true to a real-life performance environment for ecological validity.

## MATERIALS AND METHODS

### Participants

Eight pianists (4 female, 4 male) were recruited equally across classical and jazz expertise, with equivalently matched musical education to degree level. All pianists were neuro-normative and screened to be free from neurological disorders. The age range was 21 to 45. The study protocol was approved by the local Ethics Committee at Goldsmiths, University of London.

All underwent EEG recording at 1024 Hz with 64 electrodes in the 10:20 Biosemi system whilst being presented with 20 musical excerpts (10 classical and 10 jazz) via the experimental protocol below.

### Musical stimuli

A range of 20 short excerpts (4 bars or a gobbet) were presented in a visual form, i.e., the musical score and comprised a mixture of keys and tonalities (C, F, B, E flat, A flat, D flat major and E, B, F#, C# and G# minor) and tempos in a variety of rhythms (2/4, 3/4, 4/4, 6/8), motivic patterns, number of notes and melodic contours (see Supplementary Material for full table). The classical pieces (Fig 1.1) presented were unfamiliar (in consultation with syllabus assessor Richard Dawkins), and the jazz pieces (Fig 1.2) were freshly composed for the purposes of this study (composer Peter Vuust).

**Figure 1.1.**
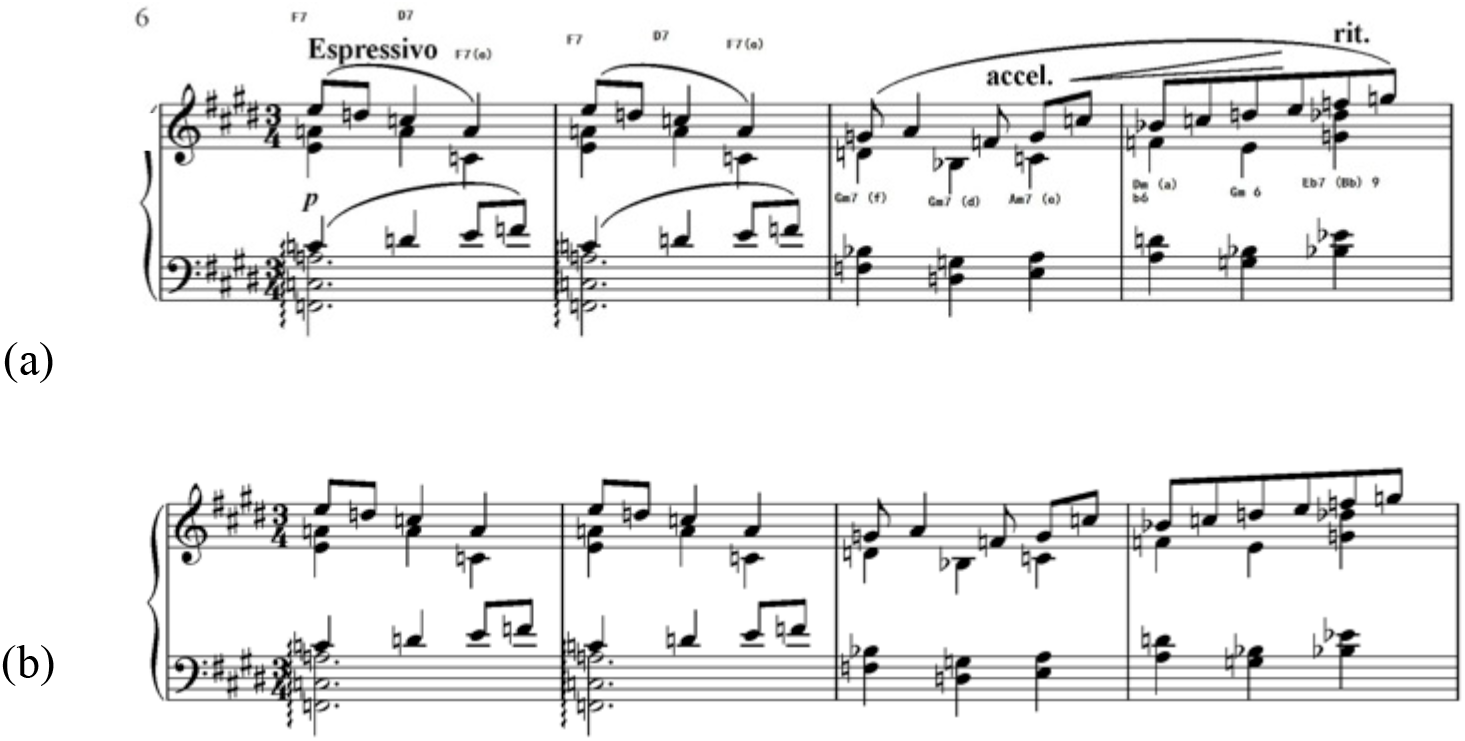
Classical excerpts with Jazz chords. (a) With expression for interpretation. (b) Without expression for play (see instruction 1) and improvisation.

**Figure 1.2.**
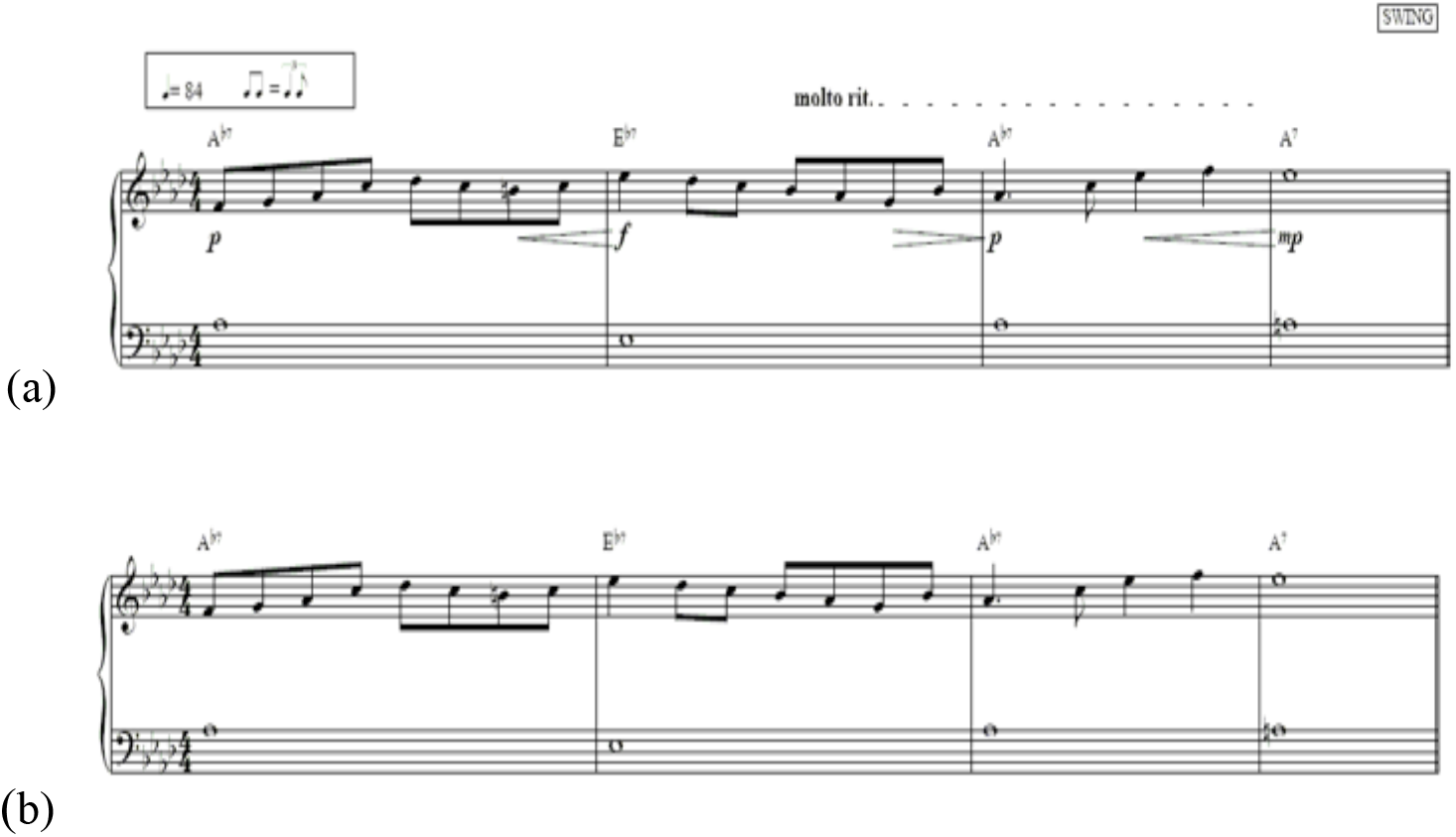
Jazz excerpts with chords and accompanying melody. (a) With expression for interpretation. (b) Without expression for play (see instruction 1) and improvisation.

### Experimental procedure

Our aim was to look at EEG brain activity during different types/stages of musical creativity and this is reflected in the experimental protocol design (please see Supplementary material for details). There were 3 separate instructions, A, ‘Play without expression’; B, ‘Interpret with expression and dynamics’; and C, ‘Improvise freely (freely improvise on the excerpt, or part thereof)’. With instruction A, musicians were asked to just play the bare notes exactly as presented, so that this instruction is considered a base resting state (verbally described to participants as “expressionless or even boring”). With instruction B, an interpretation of the composer’s markings by the musician would be a modification of tempo, emotional conveyance and volume, ornamentation, articulation and phrasing and “note inegale” such as double dotted notes, i.e., “swing” and “even eights” in jazz parlance. With instruction C, the musicians were encouraged to change the musical structure whilst still preserving a recognisable nod to the original excerpt; this would include being able to change key, melodic contour, the notes, rhythm and time signature. For each instruction, the participant was presented with just the score and given a fixed time to just think (see Supplementary Material for details) about the instruction that is presented simultaneously with the musical score and subsequently asked to physically play the excerpt in the time they would naturally take to finish without being rushed. After the completion of instruction ‘B’ (interpretation) or ‘C’ (improvisation), the participant is asked to rate their own creativity subjectively on a 5-point ratings scale: 1 being very poor, 2 being poor, 3 being ambivalent, 4 being good, 5 being excellent. Subsequently, the performances were listened to by musical judges and rated with the same scale. The order of instructions and the order of musical excerpts were randomized across participants.

There were three types of control: within subject control -when the subject is asked to play the excerpts without expression, instruction ‘A’, as a baseline; between subject control -different musicians asked to do the same protocol and perform the same classical and jazz musical excerpts; self and judges’ creativity assessments.

### Self and judges assessments: acquisition and ranking

Musical performances were rated by five judges: two from a Classical background and three from a Jazz background. The two classical judges comprised of an established professor, Julian Jacobson who is the Head of Piano from the Royal College of Music and Phil Aslangul who is the Head of Surrey Chamber Choir and is a regular assessor for the Associated Boards. The three Jazz judges were expert professional performers with high international reputation, chosen on recommendation from Simon Purcell, Head of Jazz at Trinity College Of Music. These were pianists George Fogel, Liam Noble and flautist Finn Peters.

On interview with Richard Dickins, it was indicated that a truly inspired performance was palpably ‘felt’ instantly by both performer and listener. Thence the idea arose that we should therefore ask the same question, ‘How creative did you think that was?’, to both the performing musicians as their self-evaluation and then external judges using the same ranking scaled system, based on an instant decision not subjected to prolonged over-analytical thinking. ‘Creativity’ is an integrated action so we decided to focus on the fundamental nature of it rather than employ standard creativity assessments that often measure types of thinking that could indirectly be inferred to creative thinking, such as Guildford’s different uses of a brick, or associated personality aspects such as ‘openness’, ‘emotionality’, ‘psychoticness’, etc.

Judges were sent out a marking form with the visual score of each excerpt to refer to and an audio CD with all eight participants’ performances to listen to in the same randomised order (interpret and improvise tasks), as was presented to the participants themselves. The creativity of each musical performance was rated by the judges on the same 5-point scale as the performers. They were asked to be fairly instantaneous with their “gut feeling” within the 7 second gap between each audio performance on the CD. The form used for the acquisition of judges’ ratings is included in the Supplementary Material. It includes supplementary notes to the assessors along with an example of what they were asked to do.

### EEG Setup and Signal processing

Biosemi hardware with 64 active shielded electrodes were used in the 10-20 “international spacing” standard and Actiview/Labview software was used for data viewing during acquisition. Additionally, 6 external electrodes were used, 4 ocular (2 for blinks and 2 for horizontal eye movement ‘saccades’ tracking), and 2 mastoidal (for use as reference electrodes post recording). A sample rate of 1024 Hz was set for recording with moving references whilst recording.

There was comprehensive signal processing of the raw EEG data (see Supplementary material for details). Each recording was referenced, filtered for mains electricity by a 50 Hz notch filter (IIR), high-pass filtered at 0.5 Hz (FIR) then processed by, ‘Independent Component Analysis’ (ICA) to visually pinpoint the major components alongside the scalp anatomical position contributing to each electrode signal’s wave^-^form in order to distinguish between cortical activity and ‘noise’ from artefacts such as movements, eye blinks and horizontal saccades, electric mains, malfunctioning electrodes, filters and references [Jung, 2000].

### Analysis

Source localisation was performed on the experimental EEG data collected using a software called sLORETA that allows a linear inverse mapping of the electrical activity recorded at the scalp surface onto deeper cortical structures as the source of the recorded activity (see Supplementary material for details on mechanisms). We use sLORETA as a tool on the EEG data as a detector of activity difference between different conditions and participants.

It was possible to classify the participants into their corresponding Jazz and Classical backgrounds for further sLORETA analyses based on the phenomenological interview information taken from the participants themselves. There were four in each group and comparisons were made as according to Table 1.1. When comparing the two different background groups, they were classified as independent pairs for sLORETA settings.

**Table 1.1.**
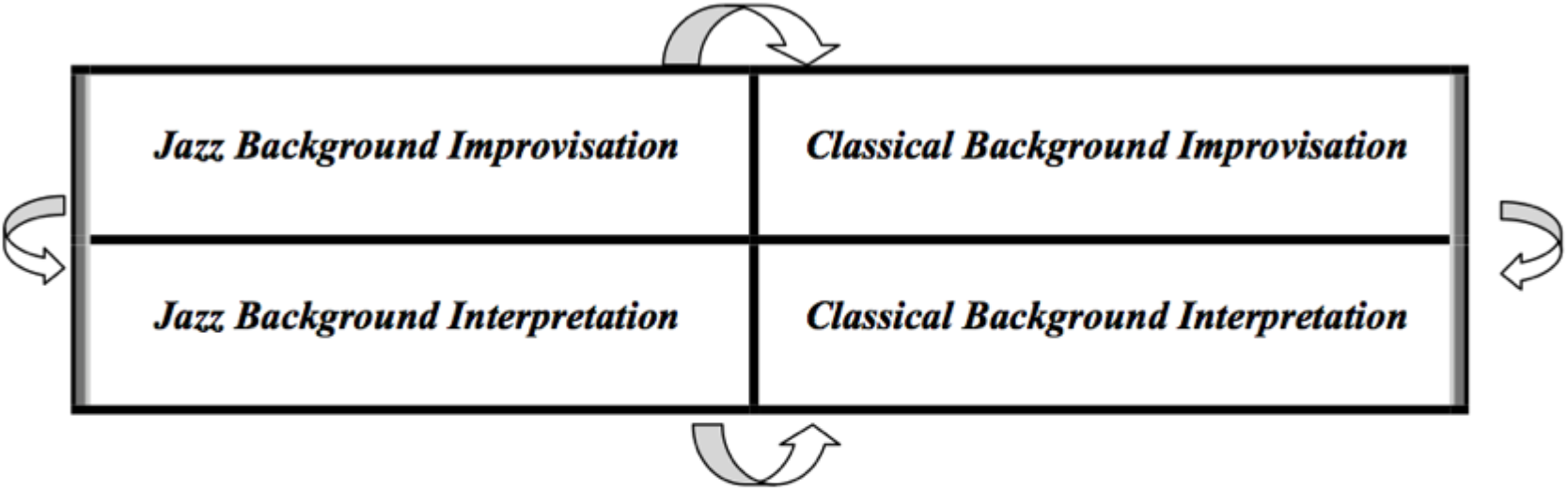
A table of comparisons made for sLORETA analysis between Jazz and Classical background participants.

For the purposes of relating assessments with sLORETA findings, we ranked the judges’ and self-assessments and used the top 78 ‘Creative’ and bottom 78 ‘NonCreative’ performances. It was noticed that although there was overall agreement between judges for individual extracts, there might be a collective bias of the Jazz judges to be systematically harsher in their assessments so this was corrected for (see Supplementary material).

Analysis of agreements between jazz and classical judges found that judges are more in agreement when they judge Jazz and Classical Interpret conditions than when it is more free such as in Improvisation (see Supplementary material for details).

## RESULTS

We summarise the major findings of the sLORETA analyses and the anatomical areas of activation that are suggested to be involved in different musical processes. We are examining the tasks ‘Interpret’ and ‘Improvise’ which are in themselves different types of creativity within music but we go a step further of what constitutes as more or less creative within these tasks.

sLORETA analyses are presented in Fig 1.3 and 1.4, on the 78 top creative and bottom non-creative extracts as based independently on judges’ mean scores and participants’ self assessments. The self-assessments and judges’ assessments were not statistically in agreement (see Supplementary material).

**Figure 1.3.**
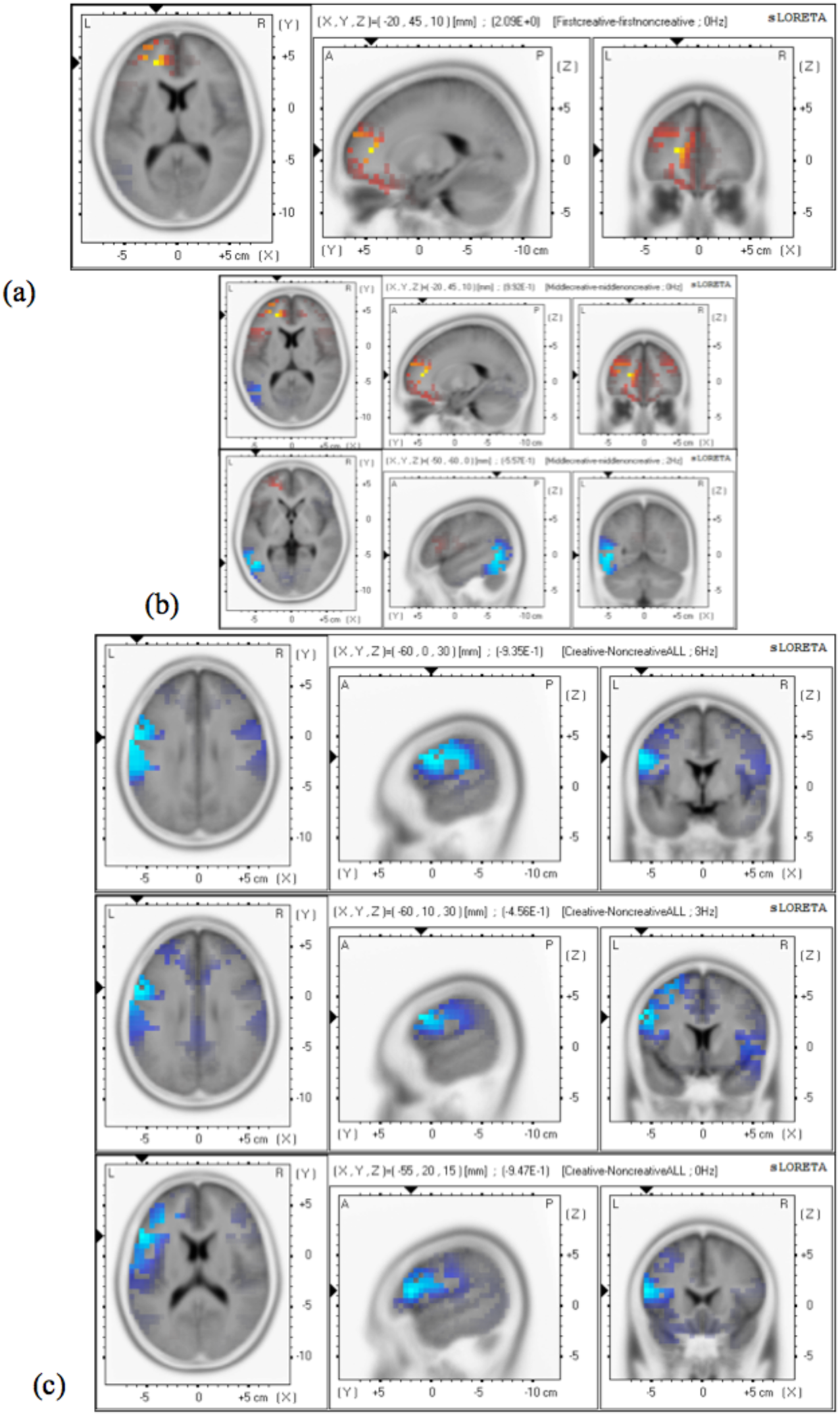
Positive and negative modulation patterns of Judge assessed comparisons of ‘Creative’ versus ‘Non-Creative’ extracts. (a) The first four seconds of performances, showing a positive modulation in the left BA 32 for the Judges. (b) The second and third rows depicts the middle four seconds of performances, showing a positive modulation in the left BA 32 and negative modulation in the left BA 37 for the Judges. (c) The last four seconds of performances, showing the negative modulations of the left hemispheric BA 6, 9 and 45 for the Judges’ dataset. Of note is that BA 32 is a robust consistent indicator of creativity in the initial and middle stages of performance using an objective external assessment. The left negative modulation of the DLPFC accompanies Jazz background creative ‘Improvisations’ versus ‘Interpretations’ as per the ‘Creative’ (80% ‘Improvisation’ tasks) and ‘Non-creative’ sets (70% ‘Interpretation’ tasks).

**Figure 1.4.**
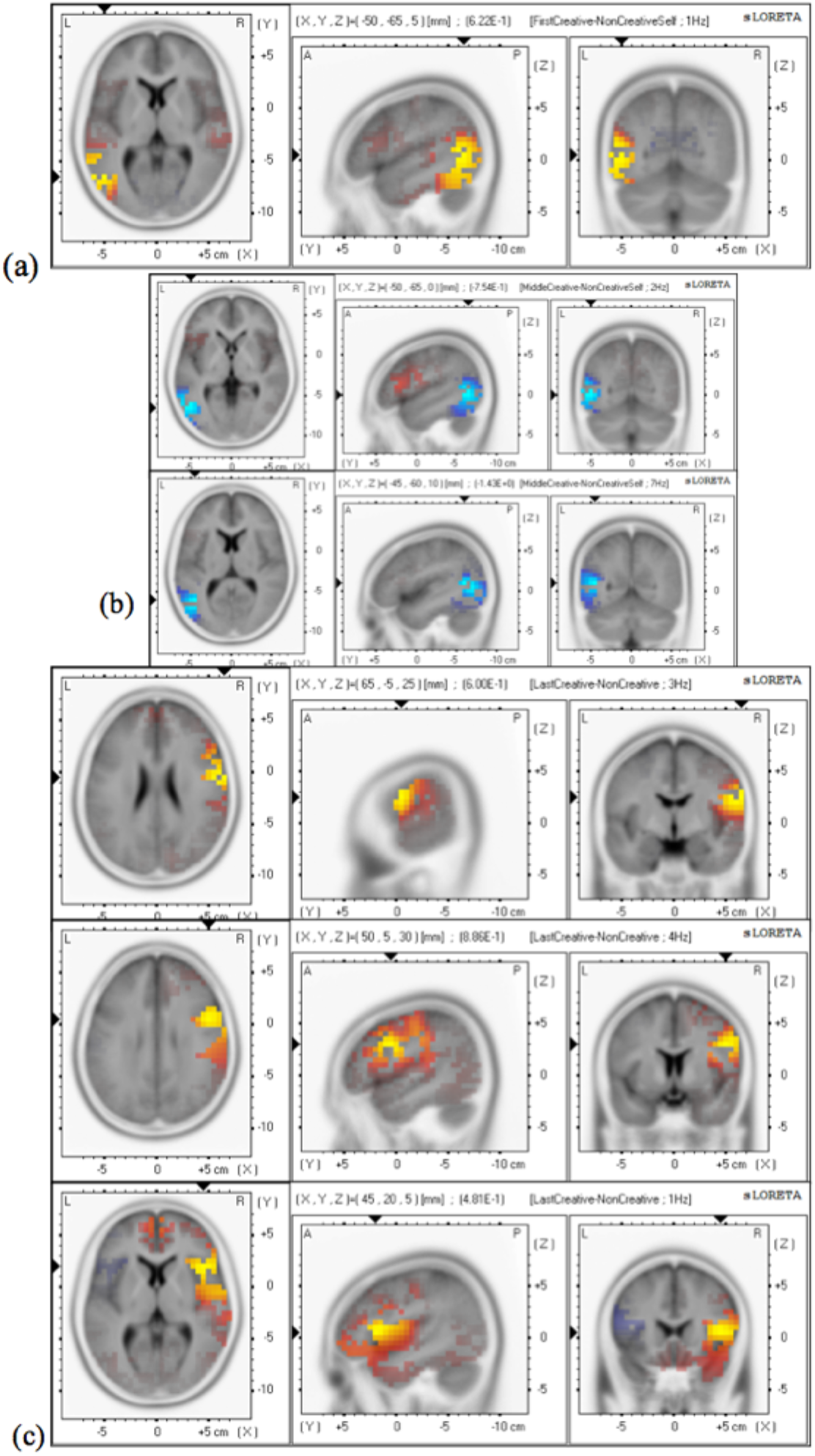
Positive and negative modulation patterns of Self-assessed comparisons of ‘Creative’ versus ‘Non-Creative’ extracts. (a) The first four seconds of performances, showing a positive modulation in the left BA 37 for Self-assessments. (b) The second and third rows depicts the middle four seconds of performances, showing a negative modulation for the left BA 37 and the left BA 39 for the Self-assessments. (c) The last 4 seconds of performances, showing the right hemispheric positive modulation in the same areas for the Self-assessments. Unlike for the Judges assessments there is no pattern that cannot be attributed to the participants’ back-ground with BA 37 to Classical and the DLPFC to Jazz. In addition, there was more fluctuation in the involved areas dependant on frequency bands, unlike BA 32 which was consistent across many frequency bands.

Using the means of judge scores, the top Creative 78 extracts across Jazz and Classical genres and ‘Improvisation’ and ‘Interpretation’ tasks were chosen between the score range between 3.78 to 5.07 and the bottom Non-Creative 78 extracts were chosen in the range 1.86 to 2.86 (Fig 1.3).

There was a consistency in the brain activity in the first and middle 4s with a positive modulation across all frequencies, of the left medial prefrontal cortex (MPFC), also known as the anterior cingulate or BA 32. In the middle 4 seconds we also see a negative modulation of the left BA 37 which we suggest is due to participants from Classical backgrounds in the sample set (see figure 1.3 and 1.6) and a persistent negative modulation pattern in the last 4 seconds of the left BA 6, 9 and 45/46 that we suggest is due to participants from Jazz backgrounds as it is only found in those with that pedagogical background.

BA 32 is suggested to be an objective indicator of creativity and is robustly present across many frequency bands in the beginning and middle 4 second sections of the ‘Creative’ minus ‘NonCreative’ comparisons for the excerpts chosen as per Judges’ assessments and not as per the self-assessments. BA 32 is not present in any other comparisons and is a likely contender to be linked to creativity, from the nature of its functional roles and anatomical links to other parts of the brain.

To examine whether this was due to (a) an objective assessment of creativity and (b) whether it was robust, we also compared extracts that were rated creative/noncreative by participants (completing the 3-way approach to this study) and three random extract sets chosen by a random number generator (see Supplementary Material for the distribution of judge means).

For the random sets, to keep the numbers of extracts consistent we simply took the top 78 Creative extracts and bottom 78 Non-Creative extracts as rated by participants themselves.

For the self-ratings, there was a positive/negative modulation pattern in BA 37 in the first and middle four seconds, similar to the temporal pattern for the tasks ‘Improvisation’ minus ‘Interpretation’ with participants from a Classical background, see figure1.4 and 1.6.

Similarly, in the last four seconds, the persistent right hemispheric positive modulation of BA 6, 9 and 45 is due to participants from Jazz backgrounds (as it is only found in those with that pedagogical background) for the same task comparison, thus suggesting an element contributable to the type of task rather than ‘Creativity’ per se. Judge and Participant ‘Creative’ datasets were both primarily ‘Improvisation’ (80% and 70% respectively), and ‘Non-Creative’ sets ‘Interpretation’ (70%), but with an overall overlap of only 40% in extracts in both sets for both ‘Creative’ and ‘Non-Creative’ extracts, this could possibly be attributed to the element of participant subjectivity, showing a specific right hemispheric positive modulation for Improvisational performances.

This overlap of 40 % could also support the positive modulation of BA 32 being due to the Judges choosing extracts that were objectively ‘Creative’ the majority of the time (i.e the other 60% of the extracts), whereas the Participants own self-assessments could be ‘hit and miss’.

The three ‘Random’ extract sets compared as such: ‘Random 1’-‘Random 2’, ‘Random 1’-‘Random 3’ and ‘Random 2’ – ‘Random 3’, showed a fluctuation of positive and negative modulations of Brodmann areas between frequencies and temporal evolutions of the performances. These were in the areas that have been mentioned before which is unsurprising as we are using extracts from an overall dataset that is from the same cognitive musical pool, but there was not a consistent pattern.

A positive modulation of BA 32 was *not* found in the Random or participants’ datasets’ comparisons.

This supports our hypothesis that there are differences in brain activity during the cognitive act of creativity. It also confirms that there is a misfit between subjective self-assessments and independent judges’ assessments and objective EEG measures consistently correlate with judges’ assessments as compared to and over self-assessments.

A positive/negative modulation pattern that emerged in the last four seconds of performance was the involvement of the insula (BA 13). It appears that there is a positive modulation in BA 13 that is specific to the condition of ‘Interpretation’ where a left hemispheric negative modulation occurs in the condition ‘Improvisation’-’Interpretation’ and a bilateral with a left hemispheric concentrated positive modulation when solely ‘Interpretation’ was compared to the baseline of ‘Play 1’, see figure 1.5. A robust pattern of positive/negative modulation appeared for BA 37, which only occurred when comparing performances from participants of a Classical Background, such that there was a left hemispheric positive modulation when comparing ‘Improvisation’ minus ‘Interpretation’ within a Classical background but a left hemispheric negative modulation when comparing the ‘Improvisation’ and ‘Interpretation’ performances between Jazz and Classical backgrounds, see figure 1.6. This suggests that for pianists of Classical background there is more use of this area possibly due to the reading from a pictorial representation of music and translating this into a musically meaningful output. They are more unfamiliar with ‘Improvisation’ performances than ‘Interpretation’ tasks and as such possibly adhere or refer back to the score more than possibly an improvising jazz musician would do. The same is true for ‘Interpretation’ tasks when comparing musicians from the two backgrounds. This is further supported by the negative modulation of BA 18 throughout any comparison of both ‘Improvisation’ and ‘Interpretation’ tasks between the Jazz and Classical participants, see figure 1.7. This area has been attributed to visual saccades [Darby et al., 1996] and also to mental imagery during music perception of pitches [Platel et al., 1997].

**Figure 1.5.**
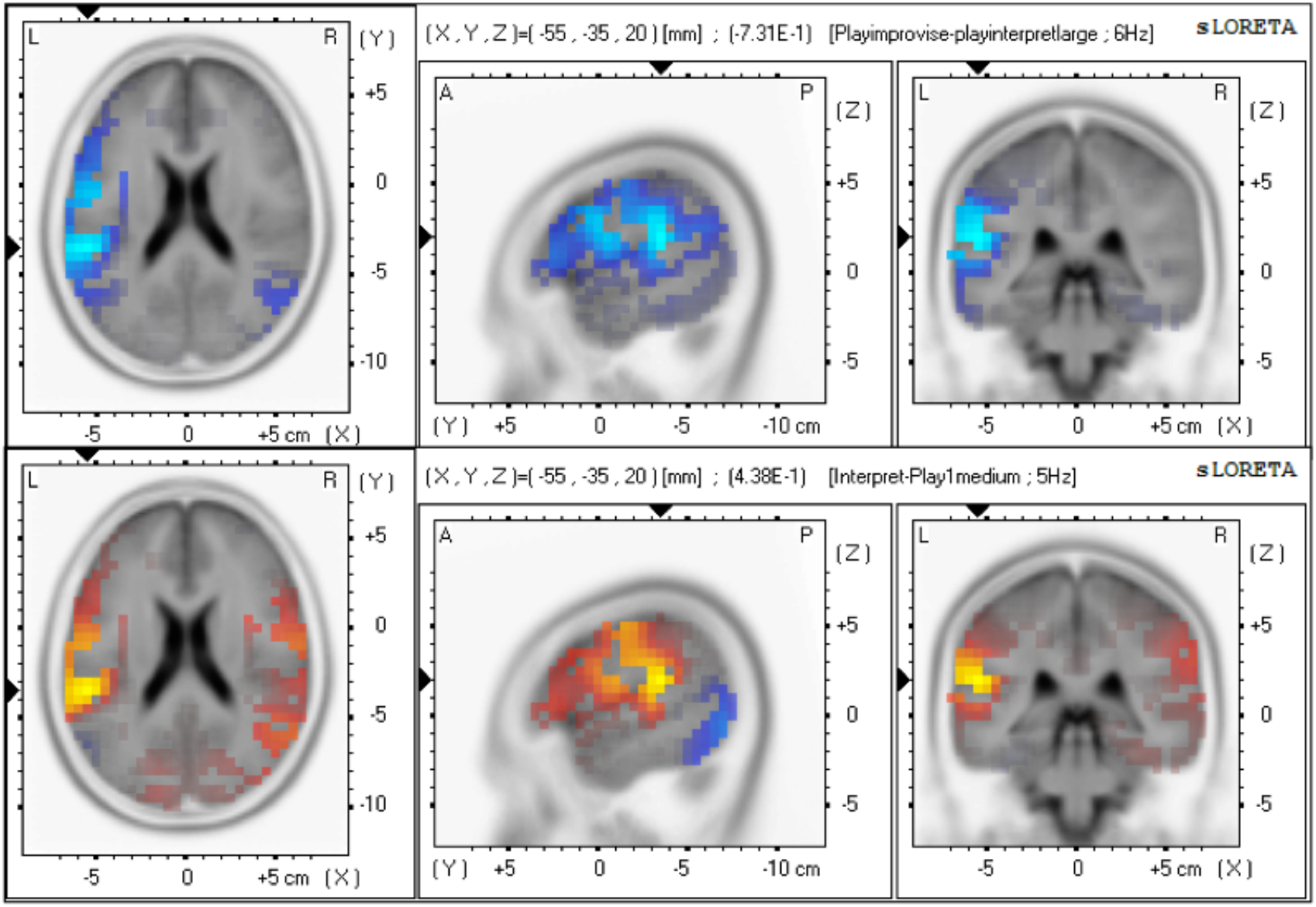
The first row depicts the comparison of ‘Improvisation’-’Interpretation’ showing a negative modulation in the left insula (BA 13) and the second row shows a positive modulation in the same area when ‘Interpretation’ is compared to the baseline ‘Play 1’. This indicates that cognitively, ‘Interpretation’ requires more conscious error-monitoring, audio-visual integration, emotionally linked response inhibition and accurate rhythmic auditory processing than ‘Improvisation’.

**Figure 1.6.**
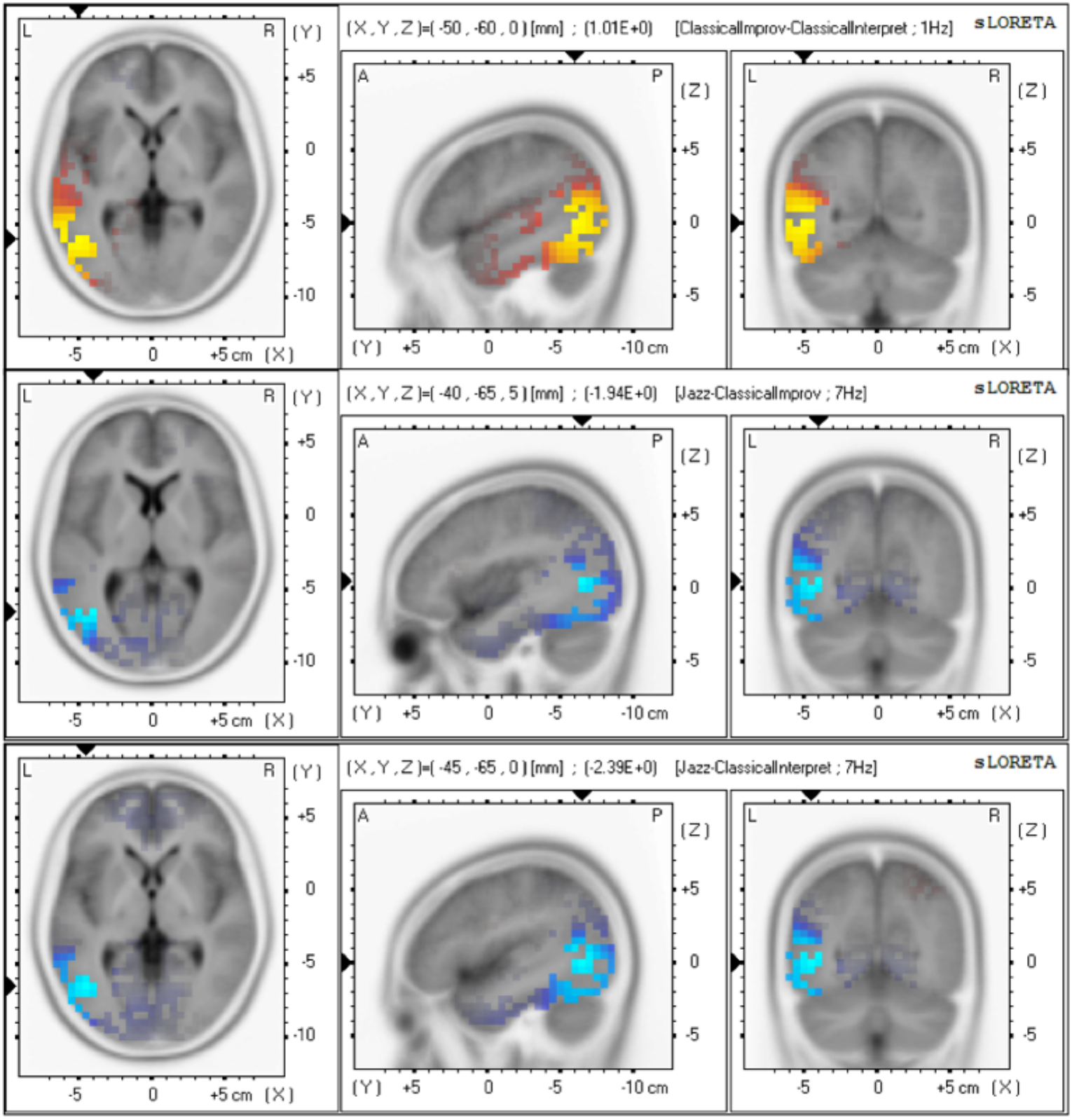
The first row depicts the condition of ‘Improvisation’-’Interpretation’ within participants from a Classical background, with a left hemispheric positive modulation of BA 37. The second and third rows depict comparisons between participants from Jazz and Classical backgrounds when comparing ‘Improvisation’ and ‘Interpretation’ tasks respectively. This indicates that the involvement of BA 37 only occurs when comparing performances from participants of a Classical background suggesting a reliance on the musical score as BA 37 is related to metaphorical/semantic processing.

**Figure 1.7.**
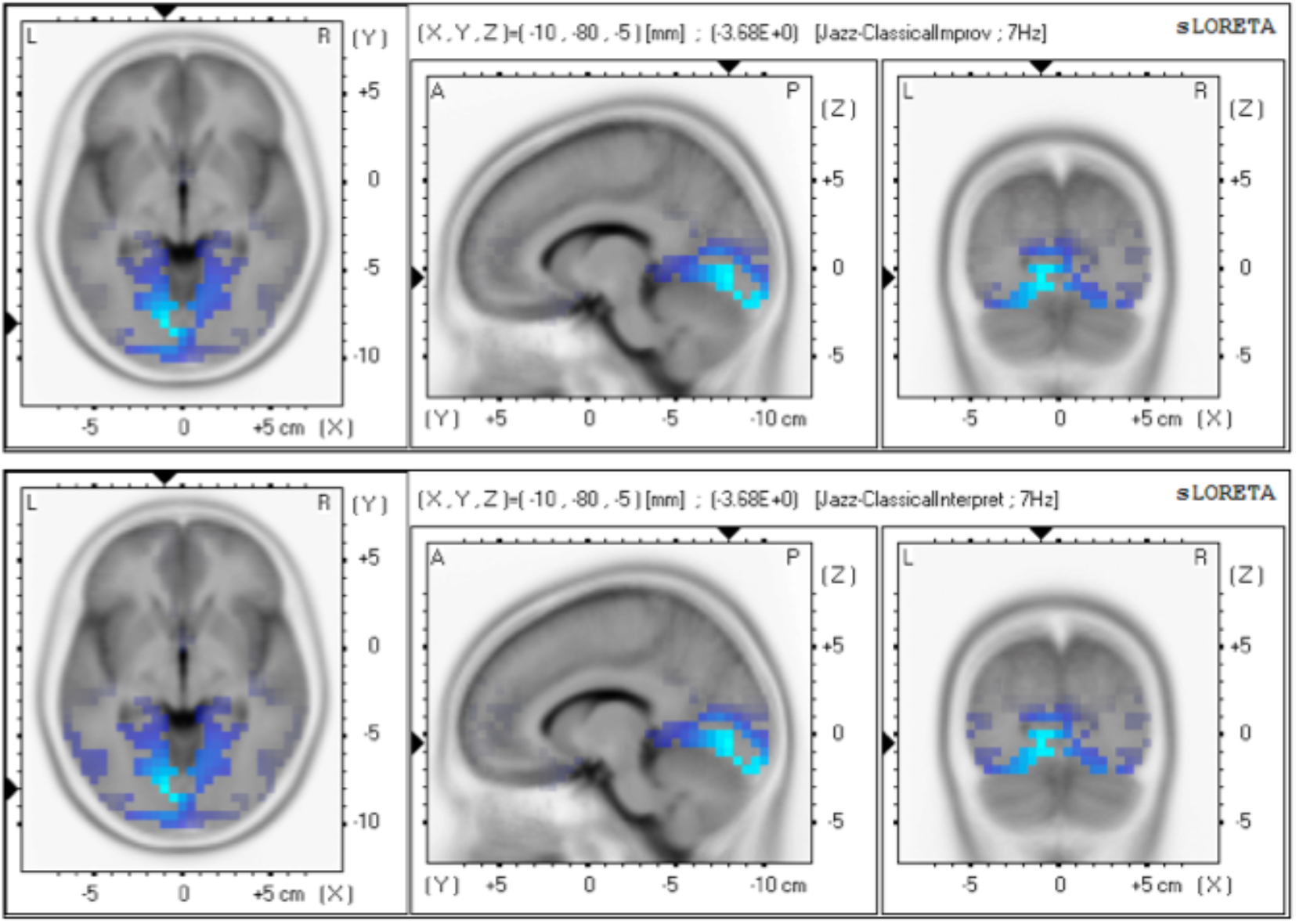
Negative modulation patterns in BA 18 for both tasks of ‘Improvisation’ and ‘Interpretation’ between participants of Jazz and Classical backgrounds. This indicates that participants from a Classical background adhere more to the visual musical score (visual saccades) and use a different form of mental imagery as compared to participants from a Jazz background.

In addition, the pattern of right hemispheric positive modulation and left negative modulation in BA 6, 9 and 45/46 is found only during ‘Improvisation’-’Interpretation’ tasks in the middle and last 4 seconds when comparing participants of Jazz background (see figure 1.8).

**Figure 1.8.**
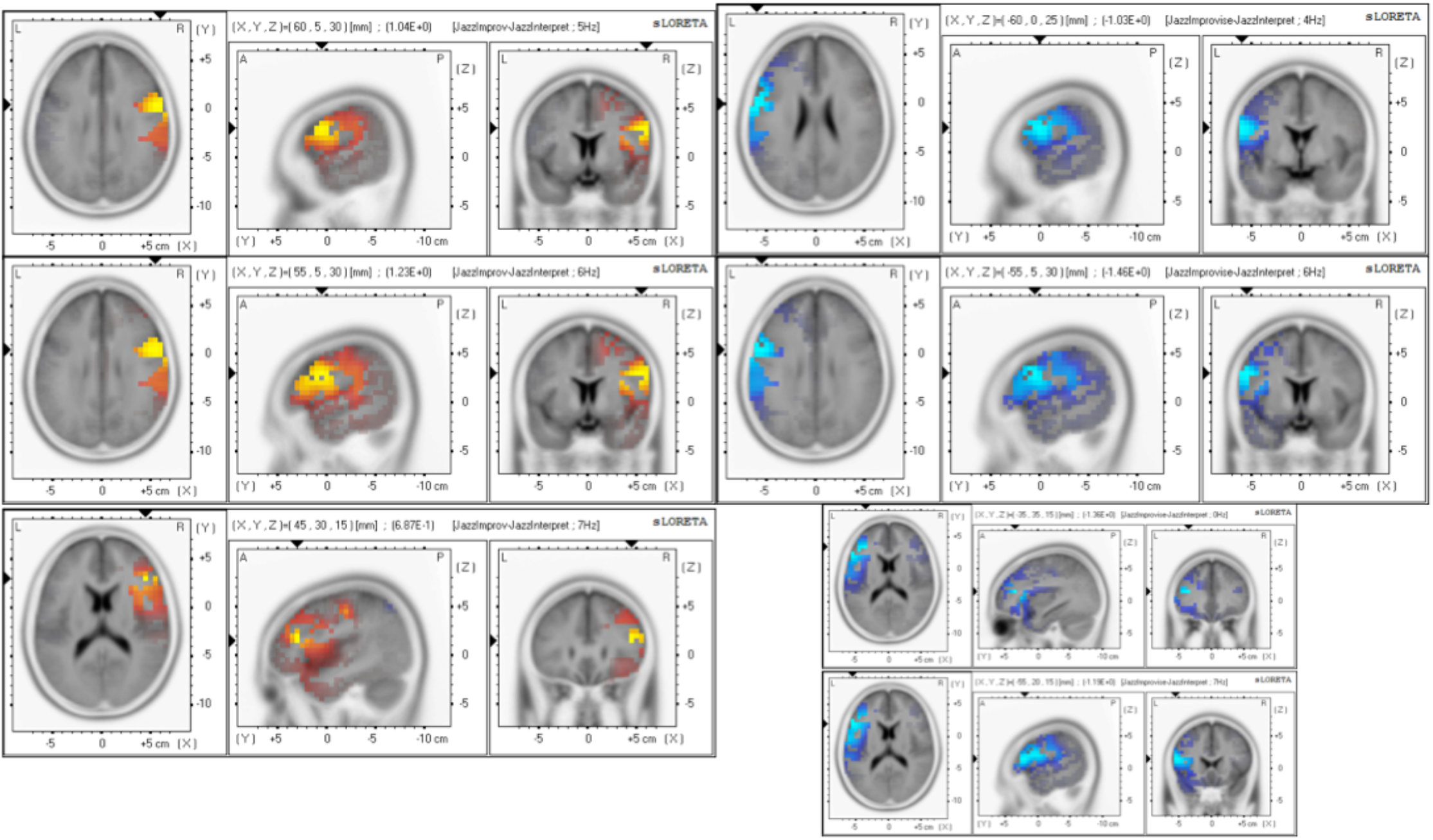
Activity patterns found when comparing the tasks ‘Improvisation’-’Interpretation’. The first row depicts the right hemispheric positive modulation of BA 6 in the middle four seconds and left hemispheric negative modulation of BA 6 in the last four seconds. The second row depicts BA 9 in the same order of positive and negative modulation, and the third row, BA 46 along with the additional negative modulation of BA 45 in the last four seconds. These are only found within participants of a Jazz background suggesting other instances of this activity pattern are due to participants’ pedagogical training.

These patterns are also found with a right hemispheric negative modulation in BA 7 in the first and middle 4 seconds again only in the participants with a jazz background for the same tasks.

This is also consistent with and is supported by the temporal pattern found during the condition of comparing ‘Improvise’ to ‘Play 1’ tasks suggesting an association specifically with the Jazz participants (see Fig 1.9). The negative modulation of BA 7 did not present itself in the condition ‘Interpret’ to Play 1’, strengthening the case that it is an area specifically related to ‘Improvisation’.

**Figure 1.9.**
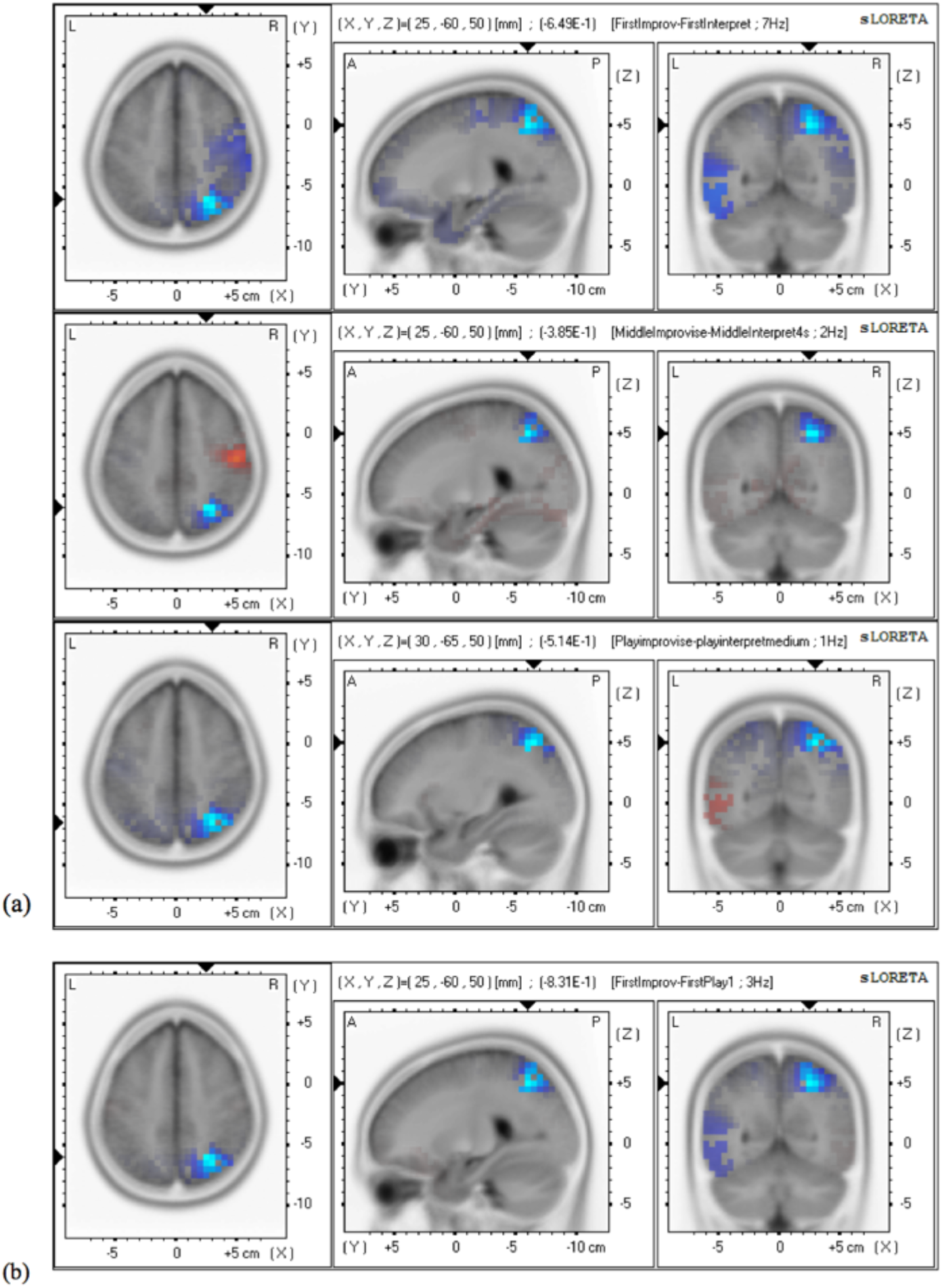
(a) Temporal evolution of the task of ‘playing’ the conditions of ‘Improvise’-’Interpret’ showing a constant negative modulation of BA 7 through beginning to end. (b) The task of ‘playing’ the condition of ‘Improvise’-’Play 1’ (baseline) also showing a negative modulation of BA 7 strengthening the possibility of its association to ‘Improvisation’. This indicates that BA 7 plays a consistent role in the task of Improvisation only rather than the other musical tasks.

The involvement of BA 7 which is also known as the precuneus or superior parietal lobe (SPL), is a constant not only in the performance tasks but when directly comparing the ‘Improvise’ minus ‘Interpret’ tasks during thinking, again there is a negative modulation of the right BA 7, alongside a negative modulation of the left BA 37, right BA 7 and a positive modulation of the left BA 40 (see figure 1.10).

**Figure 1.10.**
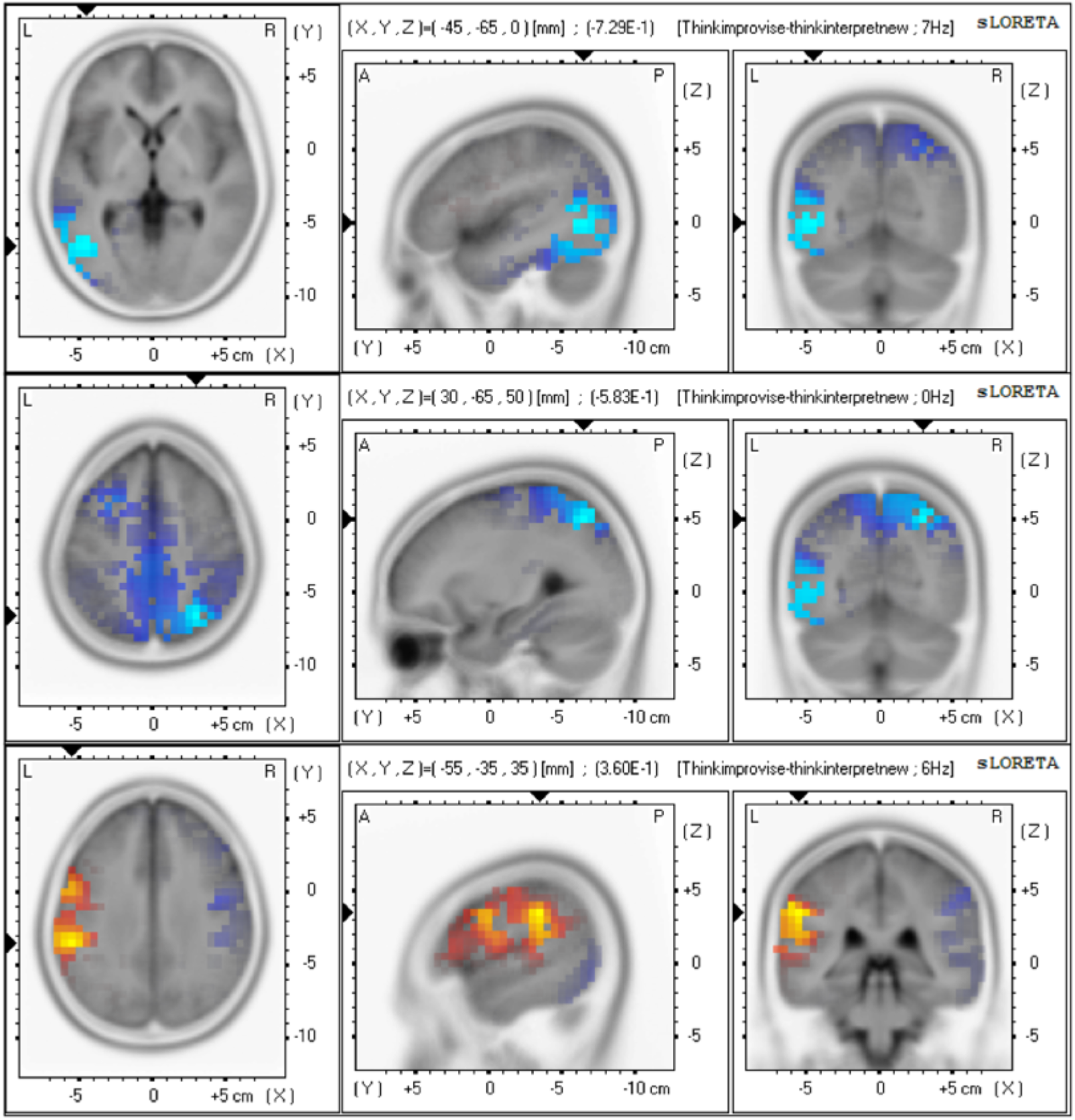
Comparing the tasks of ‘thinking’ during ‘Improvisation’ minus ‘Interpretation’ shows a negative modulation in the left BA 37, right BA 7 and a positive modulation in the left BA 40. This implies less semantic processing is required for ‘Improvisation’ and points to a different mental imagery required, in the form of an ‘Insight’ strategy. BA 7 negative modulation also associates ‘Improvisation’ with a state of consciousness that has been linked to hypnagogia, sleep-states and being under anaesthesia.

## DISCUSSION

Our results build on a study where clear differences in pre-attentive brain processing of sound feature deviations in even relatively simple musical excerpts have been recorded using EEG and MEG [Vuust 2012].

During the tasks of ‘thinking’ about ‘Improvising’, we found a distinction in the type of mental imagery used for ‘Interpretation’, which emerged in the form of Brodmann Area (BA) 40 rather than BA 37, indicating ‘Improvising’ having an ‘Insight’ strategy. This area has previously been implicated in verbal creativity for this ‘Insight’ strategy but not for music [Betchereva *et al*., 2004].

Both ‘Improvise’ and ‘Interpret’, require more of the semantic processing associated with BA 37 when compared to the baseline of ‘thinking’ about the task ‘Play’, suggesting a different form of technical musical processing with the extra cognitive load than just playing the bare bones of the excerpt presented. On closer inspection, BA 37 is an area that is specifically activated/deactivated in Classical musicians which makes sense in light of their close adherence to the score both from an educational training perspective and as a prerequisite of the task certainly when it comes to ‘Interpret’. A constant in the activity pattern for BA 37 is the deactivation in the middle segments of performance when comparing ‘Improvisation’ to ‘Interpretation’ in general and also during more creative tasks (assessed by judge and self), indicating a loss of reference to the score even with their Classical pedagogical background. The role of BA 37 in musical cognitive processing and its association specifically with musicians from a Classical background is a new finding.

BA 7 deactivation is present strongly in all Improvise comparisons during ‘thinking’ and ‘performing’ and is revealed to be something specific to Jazz musicians which also makes sense to the task from anecdotal evidence of being ‘in the zone’ as it is present during altered states of consciousness such as hypnagogia, sleep states and being under anaesthesia (see below). Previous research has linked the hypnagogic state and its corresponding frequency of theta to creativity (see section 1.4 of Rahman’s PhD thesis, 2014). An activation of BA 7, also corresponds with tasks of focussed attention and decreased bilateral activation which is linked to the integrated task of ‘Interpretation’. This area and its deactivation are suggested to be a strong feature of ‘Improvisation’ which probably uses conceptual combinations of thoughts and ideas not readily accessible at the conscious level. BA 7 also refers to less episodic memory recall and adherence to an internal mental navigation to the musical notes presented (see below). The role of BA 7 in ‘Improvisation’ amongst Jazz musicians is a new finding.

It is an area where there has not been an integrative consensus as to its function. The precuneus is very well connected with reciprocal corticortical connections in adjacent areas leading to an interconnection that is bilateral, bridging the two hemispheres and it is also selectively connected to other parietal areas such as the caudal parietal operculum, inferior and superior parietal lobules, and the IPS known to be involved in visuo-spatial information processing [Selemon and Goldman-Rakic, 1988; Cavada and Goldman-Rakic, 1989; Leichnetz, 2001].

Mainly, it is connected to the frontal lobes, namely BA 8, 9, 46, 6 and 32 [Petrides and Pandya, 1994; Goldman-Rakic, 1988; Cavada and Goldman-Rakic, 1989; Leichnetz, 2001]-the latter four of which are areas that are instrumental in the different conditions of our study.

Here we will present a few different studies that may build up a picture as to the right precuneus’ role in our study as related to mental imagery and attention. Firstly we will look at the visuo-spatial cognitive tasks that the precuneus has a role in. Relying on a lesion study, Suzuki *et al*., (1998), described a case where a haemorrhage in the right medial parietal lobe mainly in the precuneus resulted in an inability to navigate in the real world despite being able to perform well on visuo-spatial learning tests. This indicates a selective impairment of mental navigation-related networks. This is indicative of a different form of mental imagery or navigation that is internally mentally navigated, for ‘Interpretation’ tasks which is not the case in ‘Improvisation’ as indicated by the negative modulation in the right precuneus.

The SPL especially in the right hemisphere has been considered a higher order area involved in controlling spatial aspects of motor behaviour [Grafton *et al*., 1996; Connolly *et al*., 2000; Seitz and Binkofski, 2003; Grefkes *et al*., 2004], and severe disturbances of visually goal-directed hand movements (such as could be attributed to ‘Interpretation’ tasks) not related to motor, sensory, visual acuity or visual field disorders have also been ascribed to lesions of the SPL.

Specifically for music, in a PET study of musical episodic memory [Platel *et al*., 2003], melodic tunes recognition tasks contrasted with perceptive control tasks, resulted in activation of the episodic memory network, comprising the prefrontal cortex, the anterior cingulate gyrus and the precuneus. These activations were more prominent in the right hemisphere, although they were bilateral. Here, precuneus involvement was likely to be related to the success of episodic recall, as the musical material used in this experiment did not involve particularly imageable features and no subject had employed a specific mental representation strategy. This proves a likely model candidate for ‘Interpretation’ where the interpretation of composers’ markings involves recall of technique employed elsewhere and learned in training, not to mention a note-for-note reproduction from reading of score to performance. This is further supported by a PET investigation that found the left precuneus and cuneus were the main areas active during detection of pitch changes [Platel *et al*., 1997], where it was thought the pattern of activation was a consequence of the mental imagery strategy employed to perform the pitch discriminiation tasks via a ‘mental stave’.

Finally we come to the important element of attention. Hugdahl *et al*., (2000), showed that focussed attention decreased bilateral activation and an increase in the right precuneus especially for musical stimuli. This would imply that ‘Interpretation’ tasks involve more focussed attention than do ‘Improvisational’ tasks. This is further supported by a PET study on the neural correlates of visual awareness using subliminal and supraliminal verbal stimuli [Kjaer *et al*., 2001], where the right precuneus and dorsolateral prefrontal cortex were activated during visual-verbal stimulation that lasted long enough to elicit awareness. This indicated that these regions were critical for task-elicited and state-dependent awareness.

The most compelling evidence of the relationship between a negative modulation in the BA 7, attention and ‘Improvisation’ comes from a series of studies on altered states of consciousness such as slow-wave sleep (SWS), rapid eye (REM) sleep, the hypnotic state, induced-anaesthetic states and persistent vegetative states. All of these states show a profound deactivation in the BA 7 [Maquet *et al*., 1997; Maquet *et al*., 1996; Rainville *et al*., 1999; Fiset *et al*., 1999; Laureys *et al*., 1999]. It seems there is an active participation of the precuenus in conscious processes, high order body and self-representation [Maquet *et al*., 1999] which would make sense in light of the richly connected multimodal network that the precuneus belongs to. It would seem that the task of ‘Interpretation’ involves an internally guided attention and self-representation with episodic memory retrieval whilst that of ‘Improvisation’ involves more a loss of conscious self-awareness.

The activation/deactivation pattern of BA 6,9,45 and 46 from the right to left hemisphere respectively, occurs in ‘Improvisation’ minus ‘Interpretation’ comparisons only for Jazz musicians, temporally in the last segments of performance, with BA 6 specifically linked to Improvisation. Previous research has linked BA 6 to melody generation (though bilaterally) [Brown et al. 2006] and the dorsolateral prefrontal area which comprises of BA 9, 45 and 46, to lyrical creativity [Braun et al. 2012] which our research also supports by the involvement of these areas in the comparison of ‘Creative’ minus ‘NonCreative’ datasets. However, our patterns of activation are in the opposite hemisphere possibly due to a combination of the type of creativity (musical rather than verbal) and task comparison (‘Improvisation’ versus ‘Interpretation’). Our ‘Interpretation’ task seems more similar to their ‘Improvisation’ task as both are more goal-directed than our task of ‘Improvisation’. Interestingly, in the ‘Creative’ minus ‘NonCreative’ comparisons there was a constant deactivation in the left hemisphere for these areas only in the excerpts chosen with the Judges’ assessments whereas there was a constant activation in the right hemisphere only in the excerpts chosen with the Participants’ assessments. This might be a possible contender for the subjective experience versus the objective assessment in what constitutes as creative.

The activation of BA 13 or the insula during the task of ‘Interpretation’ over ‘Improvisation’, indicates a task that is more goal-orientated utilising conscious error-monitoring, emotionally related inhibition responses, audio-visual integration and rhythmic temporal auditory processing suggesting a more integrated, inhibitive, conscious and convergent creative task than ‘Improvisation’. Anatomically, the insula appears to be a candidate for multi-modal emotional and motoric integration and processing as it receives information from ‘homeostatic afferent’ sensory pathways through the thalamus and outputs to limbic areas such as the amygdala, ventral striatum, orbitofrontal cortex as well as the motor cortices [Craig, 2002]. It is well placed to be a neurobiological candidate for embodied cognition, an idea proposed by philosopher, psychologist and physician William James (1842-1910), where subjective emotional experience, i.e., conscious feelings, arise from our brain’s interpretation of bodily states that are caused by emotional events.

The association of the ‘Interpretation’ task to an increase in positive modulation in the insula could be feasibly linked to a mixture of conscious error-monitoring, emotionally linked response inhibition and audio-visual integration and temporal auditory processing as could be expected from the nature of the task.

Firstly, research by Shafritz *et al*., (2006), on an fMRI study indicated an activation in the insula in participants when inhibiting responses to emotional faces whereas a task not involving emotional valence such as during a letter task, did not strongly engage this region. Music in itself provides an emotional context to the task of ‘Interpretation’ which by sticking to the rules of the particular extract performed, is creatively more goal-oriented, convergent and more inhibitive than ‘Improvisation’.

Further research on the activation of the left insula, is a study by Klein *et al*., (2007), where activity in this region was stronger for conscious aware errors as compared to unaware errors, using an antisaccade task. What is of interest is that any post-error adjustments to do with speed and accuracy in performance was only observed after aware errors which is in keeping with the nature of ‘Interpreting’ a piece of music where accurately adhering to the notes and dynamic instructions is of utmost importance and requires a conscious awareness during error-monitoring.

A cross-modal study by Lewis *et al*., (2000) involving the perception of speed of an auditory signal and the simultaneous identification of visual dots with the highest velocity, showed an enhancement of activation in the left insula. Lewis et al. theorised that this polymodal effect could have reflected specific task factors, such as attentional tracking of the target, selection/computation of the relevant motion parameter (speed), comparison of speeds, selection of response, or non specific task factors such as storage and retrieval of information from working memory, all of which are cogntive tasks collectively involved in the overall musical task of ‘Interpretation’. The task of ‘Interpretation’ requires a greater integration of information from the visual cues of the musical score and the auditory sensory perception and monitoring of the actual music played, requiring an essential audio-visual integration that is not as necessary in the task of ‘Improvisation’, where participants are only required to use the visually presented musical score as a starting point for their performance and not as a continual reference required throughout their improvised performance. Participants could, if they wanted to, improvise with their eyes closed, whereas unless they memorised the extract in the few seconds it was presented for, they could not do so whilst interpreting it.

Finally, the left insula has been shown to be important in perceiving the regularity or irregularity of a temporal rhythm within a musical sequence [Platel *et al*.,1997]. The authors attributed the role of the left insula to memory processing whereas Colavita *et al*., (1974), postulated that the insular-temporal region is crucial for discriminating temporal auditory patterns by attending to the entire pattern as a whole through their work in cats. Interpreting a piece of music certainly requires an awareness of the whole shape of the piece an an ability to temporally hold in the mind what has just been played, what is being played presently and preparing for what is about to be played in the future. It also requires an accurate knowledge of different time signatures and rhythms as indicated by the composer and the ability to carry this out with accuracy. Again in ‘Improvisation’, this is less externally goal-oriented and more an internal choice or preference, with the ability to change rhythms, tempi and time signatures at will and ‘in their own time’ if you will, whereas unless indicated by the composer, this is not the case with ‘Interpretation’.

The involvement of the insula in the cognitive musical task of ‘Interpretation’ indicates that it is one which requires conscious awareness, emotionally related inhibition responses, error-monitoring, audio-visual integration and rhythmic temporal auditory processing suggesting a more integrated, inhibitive, conscious and convergent creative task than ‘Improvisation’.

Most importantly, BA 32 is suggested to be an objective indicator of creativity and is robustly present across many frequency bands in the beginning and middle sections of the ‘Creative’ minus ‘NonCreative’ comparisons for the excerpts chosen as per Judges’ assessments and not as per the self-assessments. BA 32 is not present in any other comparisons and is a likely contender to be linked to creativity, from the nature of its functional roles and anatomical links to other parts of the brain. It allows an integration of motoric and emotional communication with a maintenance of executive control, consciously monitoring and implementing adjustments in an ongoing performance. This positive modulation of the left MPFC supported the findings from Braun *et al*., (2012) for their ‘Improvisation’ versus ‘Conventional’ comparisons and makes for a good candidate for creative performances when compared to those that are non-creative, as the area has many afferent and efferent connections. It allows a maintenance of executive control, consciously monitoring and implementing adjustments in an ongoing performance [Miller and Cohen, 2001; Tanji and Hoshi, 2008].

This finding supports previous research in lyrical verbal creativity, but is novel in musical creativity also because it suggests that judges make their decision for a highly creative performance early within hearing a performance and this is also reflected in the EEG of the performers themselves. This suggests that performances that are more creative are more coordinated from the beginning and is supported by recent phenomenological research presented by Lubart and Botella (2013, MIC Conference) on the creative processes of art students characterised from interview and working-diary analyses. Functionally, BA 32 seems to be a good contender as an early indicator as it has in previous research been seen to operate at the surface of intention and action, synthesising information, encoding goals and guiding self-generated, stimulus-independent behaviours. The early recognition by both assessors and the participants (indicated by their EEG pattern of activation) lends weight to the original premise behind our choice of creative assessment in trusting the ability of both performer and audience to know instantly if an inspired performance was to ensue, by simply asking both participants and judges, the question of ‘How creative did you think that was?’. Furthermore, the MPFC has been linked as a neurological correlate in the ‘Valuation Circuitry’ during economic decision-making when integrating the various dimensions of an option into a single measure of its idiosyncratic subjective value to then choose the option that is most valuable [Kable and Glimcher, 2009]

This association between the judges’ assessments and EEG completes a loop by confirming an objective psychological impression with neurobiological evidence which could act as an early biomarker in the investigation of ecologically valid musical creativity and is a novel finding. Using sLORETA, we have been able to highlight a few anatomical areas that functionally contribute to the musical brain.

## CONCLUSION

The study of creativity is an important challenge for complexity science. It requires a breadth and depth of understanding that needs to be reflected in the comprehensiveness of any experimental design of research done into it. Coloured by as many disciplines and genres that can manifest creativity, there is also the question of the many branches of academia that pursue it, from neuroscience to musicology and philosophy to mathematics, creativity rests under the heading of a complex system as a truly interdisciplinary endeavour. Our challenge has been to integrate the many avenues and approaches into an ecologically valid study of musical creativity in particular. Source localisation was performed on the experimental EEG data collected using a software called sLORETA that allows a linear inverse mapping of the electrical activity recorded at the scalp surface onto deeper cortical structures as the source of the recorded activity. Brodman Area (BA) 37 which has previously been linked to semantic processing, was robustly related to participants from a Classical background and BA 7 which has previously been linked to altered states of consciousness such as hypnagogia and sleep, was robustly related to participants from a Jazz background whilst Improvising.

In order to characterise a topic that is in danger of almost being too vast to contain or even conceptualise, we thought to pinpoint a few key markers that are both creativity’s drivers and its manifestations, by in the first instance attempting to quantitatively note agreements between a generator and an observer of creativity. These agreements are inherently self-referential inside the creative system by its very nature, and thus both participants and external judges were given the same assessment ratings of 1 through to 5 (1 being very poor and 5 being excellent), and asked the same question based on a gut instinct assumed to be felt by both, of ‘How creative did you think that was?’.

As predicted, the self-assessments and judges’ assessments were not statistically in agreement. Self-assessments also had a lack of consistency as compared to judges’ assessments. Furthermore, based on the ratings alone, it was possible to classify participants into either Jazz or Classical, which agreed with phenomenological interview information taken from the participants themselves.

Which brings us to the third key marker of detecting a signature with no prior assumption of the underlying neurobiological mechanism but a dearth of qualitative and some quantitative observational knowledge from other researchers.

By investigating extracts that were deemed to be creative by independent judge assessments, we completed a loop, confirming an objective psychological impression with the neurobiological evidence of source localisation activity in the left hemispheric Brodmann Area (BA) 32 as an early and robust indicator of creativity during a performance within the performer’s brain. This was a novel indication in the context of musical creativity and was supported by research into verbal lyrical creativity. Strategically placed, it is an area that is particularly well connected and allows an integration of motoric and emotional communication with a maintenance of executive control.

BA 37 which has previously been linked to semantic processing, was robustly related to participants from a Classical background.

BA 7 which has previously been linked to altered states of consciousness such as hypnagogia and sleep, was robustly related to and consistently deactivated during both thinking and performing ‘Improvisation’ by Jazz musicians. This was another novel result and this area has previously been linked to a default ‘wandering mind’ network; this could suggest that the sub-process of ‘Incubation’ contributed to ‘Improvisation’ and that it is a divergent creative task.

Across genre backgrounds, within the performance phase for the interpretation task compared to the improvisation task, there was an increased activity in the insula (BA 13), suggesting a convergent creative task from the linked goal-orientated conscious error-monitoring and audio-visual integration functions.

Conventionally speaking, Jazz musicians usually improvise and Classical musicians interpret, but in our study we have placed them out of their comfort zones by asking them to do the other musicians’ normative tasks. However, we can infer from the results that the type of musical creative task is closely linked to the genre of music as can be seen by the activation of BA 7 only for Jazz musicians whilst improvising which favours lower states of attention and divergent thinking processes, the activation of BA 13 only during interpretation which favours conscious error-monitoring and integrated convergent thinking processes, and the activation of BA 37 only for Classical musicians that underlies semantic processing and a close adherence to the musical score.

### Outlooks

The presence of the left BA 32 in extracts deemed to be objectively creative by external judges, indicates a role of integration in the initial stages of a creative task-a proverbial ‘deep breath’ before starting, which conceptually fits expectations. BA 32 being anatomically well-connected, indicates that it acts as an important hub in the musical brain network that allows access to disparate areas and therefore possibly subsystems. Therefore, the directionality and temporal characteristics of BA 32 connectivity should be further investigated with network analyses for global state and temporal complexity characterisations, using Granger Causality and directed Mutual Information. Having seen a temporal evolution in activations within the different types of performances, it would certainly be worthwhile to investigate temporal network dynamics in more detail now that we have localised the key functional areas of BA 32, 7, 13, and 37.

## For the acknowledgement

Richard Dickins, Avgoustos Psillas, Matthew Lee Knowles, Simon Purcell, Julian Jackobson, Finn Peters, Philip Aslangul, Liam Noble and George Fogel,

