## Supplementary Material for "Neural markers for Musical Creativity in Jazz Improvisation and Classical Interpretation"

| Time Signature | |
|---|---|
| 4/4 | 5 |
| 2/4 | 4 |
| 3/4 | 2 |
| 5/4 | |
| 6/8 | 3 |
| 3/8 | 2 |
| 12/8 | 1 |
| 4/8 | 1 |
| 2/2 | 1 |
| 6/4 or 3/2 | 1 |
| 9/8 | 1 |
| 5/8 | |

| Number of Flats/Keys | |
|---|---|
| None (sharps or flats)/ C Major | 3 |
| One / F Major or D Minor | 2 |
| Two/ B Major or G Minor | 2 |
| Three/ E flat Major or C Minor | 1 |
| Four/ A flat Major or F Minor | |
| Five/ D flat Major or C Minor | 1 |

| Number of Sharps/Keys | |
|---|---|
| One/ G Major or E Minor | 3 |
| Two/ D Major or B Minor | 4 |
| Three/ A Major or F# Minor | 1 |
| Four/ E Major/C# Minor | 3 |
| Five/ B Major/G# Minor | 1 |

**Table 1.1.** A table displaying the tally of the extracts (20 experiment and 1 practice) that contained different time signatures and keys.

**Experimental protocol**

All participants were presented excerpts in the same settings with the same instructions on a computer screen. Excerpt and instruction presentations were automated and the timings of these and any midi information from the keyboard were superimposed onto the EEG recordings simultaneously as markers. This was so that it would be a very accurate time division of the different tasks, during analysis of the EEG.

There were 3 different instructions when an excerpt was presented such that the order of instruction was ‘A’, either ‘B’ or ‘C’ (randomized), then ‘A’ as a return to base line, and once more either ‘B’ or

‘C’ (randomized). [‘A’ is ‘Play without expression’, ’B’ is ‘Interpret with expression and dynamics’, and ’C’ is ‘Improvise freely (freely improvise on the excerpt, or part thereof)’].

For each instruction, the participant was presented with just the score for 4 seconds and then given fixed times to just **think** (11 seconds for instruction ‘A’ and 16 seconds for instructions ‘B’ and ‘C') about the instruction that is presented simultaneously with the musical score and subsequently asked to physically play the excerpt in the time they would naturally take to finish without being rushed.

After the completion of instruction ‘B’ or ‘C’ the participant is asked to rate their own creativity subjectively via pressing 1 of 5 keys in the lowest octave on the keyboard itself that correspond to a rating. Participants were put through a practice run with two excerpts that were not included in the actual experimental run.

**Judges' Form (and CD/mp3s plus data on server)**

Dear Assessor,
Thank you for agreeing to take part in this research. Your participation is invaluable to the success of this innovative research. When we have concluded the project we would be delighted to invite you to a presentation of how your contribution fits into the wider context of our project.

The task is to assess the performances that you will hear on the accompanying audio.

The score for each extract (either classical or jazz) will be presented visually to you but you will hear each extract being played as per two instructions given to the subject:

1. '*Please interpret this extract using the composer's markings included'*
   (shortened in the tables below as "Interpreting")
2. '*Freely improvise on this extract (or a part thereof)'*
   (shortened in the tables below as "Improvising")

For "freely improvise", the participants were told that there had to be something audially recognisable that linked their improvisation to the original extract. The order of these instructions are randomised in the experiment and you will hear a voice clearly indicating the sections as either "Interpreting" or "Improvising".

A voice will then ask you to rate the creativity of the subject either interpreting or improvising - you will be given a moment to do this,alternatively you may wish to pause if longer needed. Please rate creativity on a scale of 1(very uncreative) to 5 (very creative) by ticking the appropriate box as indicated by the following example table:

**Extract 1:**

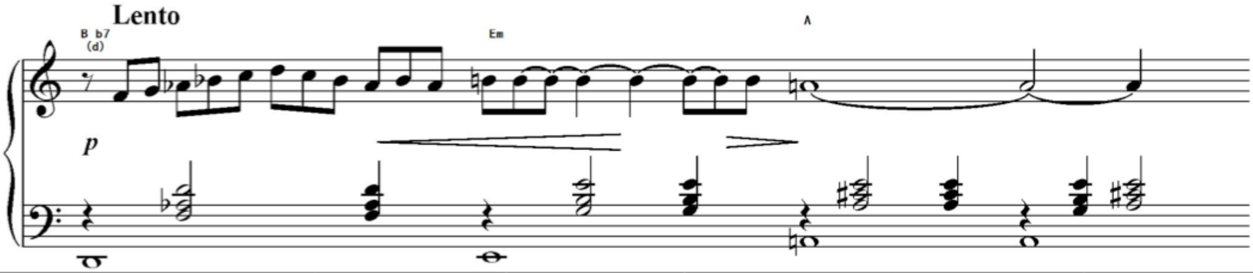


| **Creativity ratings** *Was the subject creative when* | **1** v. uncreative | **2** uncreative | **3** ambivalent/neutral | **4** creative | **5** v.creative |
|---|---|---|---|---|---|
| **Improvising** | √ | | | | |
| **Interpreting** | | | | √ | |

Please attempt to complete one participant assessment per sitting (approx 20 min) in order to maintain consistency of assessment. Please be aware that external stimulants/factors i.e. coffee/alcohol, time of day could have a significant effect on your perception/partiality.

Note that the emphasis of this assessment is on creativity rather than accuracy and that the participants are from a diverse background ranging from classical and contemporary to jazz. All have sight-reading ability but will tend to interpret and improvise on the score according to their education.

Thank you very much for your time,

Shama Rahman

**EEG Equipment and Setup**

Here we summarise the standard EEG equipment, procedure and setup used. Biosemi hardware with 64 active shielded electrodes were used in the 10-20 “international spacing” standard and Actiview/Labview software was used for data viewing during acquisition. Active shielding minimises noise from mains electricity though it is not so protective against movement artefacts hence participants were asked to keep their heads as still as possible though there were no restrictions with their hands. Participants were fitted with the appropriately sized scalp hat and their skin abraded for the placing of the electrodes which were then individually fitted and skin resistance checked until optimal. Additionally, 6 external electrodes were used, 4 ocular (2 for vertical blinks placed above and below the right eye, and 2 for horizontal eye movement ‘saccades’ tracking, placed at the outer canthus of each eye), and 2 mastoidal (for use as reference electrodes post recording). A sample rate of 1024 Hz was set for recording with moving references whilst recording so that reference could be set post recording. EEG signals were algebraically re-referenced against the average of two mastoid electrodes.

**Signal Processing**

There was a detailed and comprehensive processing of the raw EEG data. This involved EEG data recorded for each phase, task and excerpt for 8 participants leading to a total of 1,280 separate EEG recordings to process and clean before analysis.

Each recording was referenced using a GUI called EEGLab. Systems with active electrodes (e.g. Biosemi Active Two), record data “reference-free". A "zero" reference must be chosen *post hoc* during the data import and failure to do so leaves 40 dB of unnecessary noise in the data. The reference can be relatively more neutral places (i.e. with less electrical activity than the scalp electrodes of choice) such as the mastoids, earlobes or nose. In our experiments, the external electrodes were placed on the mastoids.

Again with EEGLab, each recording was then filtered for mains electricity by a 50 Hz notch filter (IIR) and bandpass filtered above 0.5 Hz (FIR) before being concatenated into one EEG recording per subject for the purposes of running a machine-learning algorithm, ‘Independent Component Analysis’ (ICA). This helps visually pinpoint the major components contributing to each electrode signal’s waveform and each component can be localised to a position anatomically on a scalp map. If there are 64 electrodes plus 6 external electrodes and 2 status channels, the ICA will return 72 components. It is based on the Principal Component Analysis and the EEG ‘cleaning’ is analogous to solving a problem similar to the ‘Cocktail Party’ problem: what a guest finally hears is a blend of sounds coming from several sources in the room i.e. the host, the other guests, several music speakers dotted around. The algorithm allows us to pinpoint these individual separate sound sources using only the merged ‘blend’ of the final sound which is similar to the ‘cocktail’ of EEG signals we record at the scalp where signals will be a mixture of cortical activity and ‘noise’ from artefacts such as movements, eye blinks and horizontal saccades, electric mains, malfunctioning electrodes, filters and references. Consequently, some of these components can be different from actual cognitive activity that we are interested in and can be rejected individually by eye, by carefully considering their waveforms and corresponding power spectra.

Electrical interference of 50 Hz, or electrical artefact bursts caused by clenching of the jaw/shoulder muscles or blinking of the eye, permeate throughout the EEG recordings of this cognitive activity and result in generally ‘noisy’ recordings. Fortunately, the movement-related ‘noisy’ electrical activity can be sourced from the 6 external electrodes as they are positioned near the eyes and jaws. The algorithm thus uses these external electrodes as a reference to separate and discard, unwanted ‘noisy’ components from the cognitive components present in all of the 64 electrode recordings. ICA can therefore most commonly detect very strong pulses such as eye blinks and horizontal eye saccades that are a necessary by-product of an experiment that relies on reading music.

ICA is also able to detect filter and mastoidal reference artefacts and due to the reliance on electrical equipment, any residual 50Hz activity that appears as interference in electrode activity even after the notch filter is applied. When the ICA waveform for this 50Hz component is examined, it looks ‘noisy’ with an increased frequency and amplitude and its corresponding power spectra does not have a clean peak at 50 Hz.

ICA is also able to detect generally malfunctioning noisy electrodes, or that are ‘drifting’ due to a loose connection, that are still present and ‘noisy’ even after being corrected for by the 0.5Hz band pass filter. The ICA algorithm commonly returns the top 10 components in order first and non-surprisingly, they consist of eye movements, filter and reference artefacts, electrical interference and noisy/malfunctioning/drifting electrodes whilst movement artefacts are finer and more spread out amongst the other remaining components. After the automation of filters, referencing and the running of the ICA algorithm, all these artefacts have to be removed individually by eye.

The underlying mechanisms of ICA, how it relates to the more commonly known Principal Component Analysis (PCA) along with a more detailed explanation of how ICA components make up a recorded waveform and why it can be used for artefact rejection can be found in a paper by Jung, Makeig and colleagues, Makeig being one of the creators of the freeware EEGLab that allows us to perform ICA and reject components [Jung, 2000].

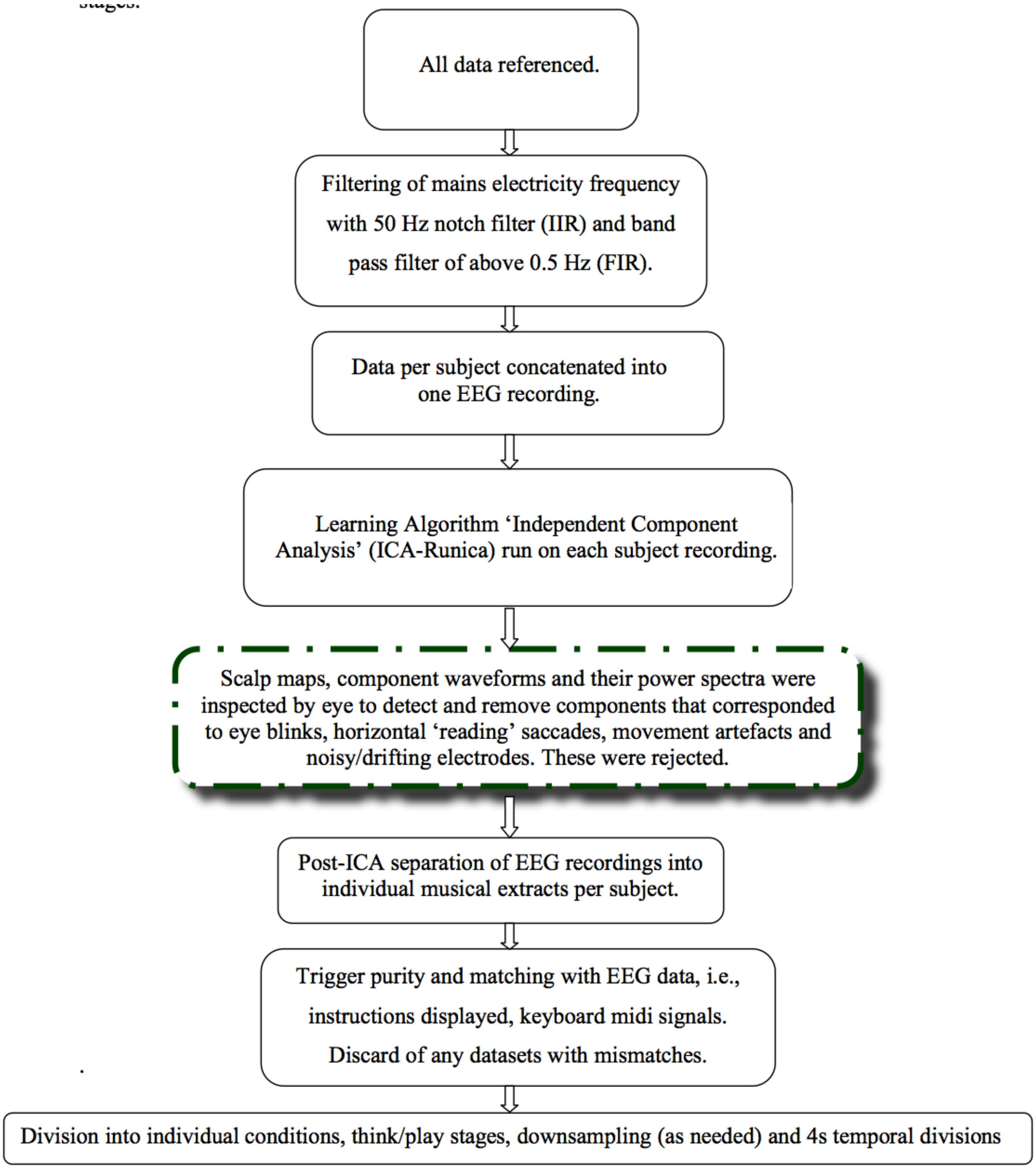


**Fig 1.1.** A flow diagram of the steps required in the signal processing of each EEG recording. The Stage highlighted in green will be elaborated in further detail below.

**sLORETA Mechanisms**

sLORETA is a functional brain imaging method that uses a quantitative neuroanatomical digitised Talaraich atlas of the cortical structures in the brain provided by the Brain Imaging Centre, Montreal Neurological Institute. The cortex can be modelled as a collection of volume elements (voxels) in this digitized Talairach atlas similar to the units found in fMRI.

As was mentioned above, fMRI and other functional imaging methods such as PET (Positron Emission Tomography) are limited in their temporal resolution and not sufficient for the speed at which neuronal processes occur. A study by Logothetis *et al*. (2001), showed that the time course of the fMRI haemodynamic response was roughly a low pass filtered (i.e., low time resolution) version of the electric neuronal activity.

On the other hand, EEG/MEG surface scalp measurements do not contain sufficient information on the three-dimensional (3D) distribution of electric neuronal activity for deeper cortical structures as the implication is that the extracranial measurements could be due to many different distributions of cortical electrical generators [Helmhotz, 1853] known as the inverse problem.

Ideally, it would be optimum to utilise both the temporal resolution afforded by experimentally recorded extracranial signals and localise the brain activity source of these signals by solving the inverse problem. Given that brain activity occurs in the form of a finite number of distributed “hot spots”, using the principles of linearity and superposition, would allow the calculation of an instantaneous, distributed, discrete, linear solution capable of exact localization of point sources  [Pasqual-Marqui, 2002].

**Judges' assessments: Corrections**

It was noticed that although there was overall agreement between judges for individual extracts, there might be a collective bias of the Jazz judges to be systematically harsher in their assessments.

Two Centres of Mass (CM) for the three Jazz judges and the two Classical judges, were calculated separately for the Classical and Jazz extracts, leading to a total of four values: CM(JJ), CM(CJ), CM(JC),CM(CC), where JJ depicts Jazz Judges' assessments for Jazz extracts, CJ depicts Classical Judges for Jazz extracts, JC depicts Jazz judges for Classical extracts and CC depicts Classical judges for Classical extracts, e.g.

$$CM(JJ) = \frac{\sum_{JJ} scores}{N_{JJ}}$$

where $N_{JJ}$ is the total number of JJ assessments. These were then plotted as two co-ordinates (CM(JJ), CM(CJ)) and (CM(JC), CM(CC)), depicted by the stars in Figure 24 'a' and 'c' superimposed on the Classical and Jazz Judge scores for the separate batches of Classical and Jazz extracts. The deviance from the identity line (y=x), confirms that the Jazz judges were indeed marking harsher for both Jazz and Classical extracts.

The deviances were then corrected for in the Jazz judges in the separate batches of Jazz and Classical extracts by adding the difference from the CM of the Classical judges to the individual Jazz judges' marks for each extract i.e., 'CM(CJ) - CM(JJ)' for the Jazz extracts and 'CM(CC) - CM(JC)' for the Classical extracts, see Figure 5.2.

After this correction, a consistency is achieved between the different types of judges so that the assessments are calibrated and means are equal to each other. From here on we will use the corrected Jazz judge assessments together with the Classical Judges assessments, for a more accurate representation of the external evaluation of the participants' performances. As such it is these calibrated corrected means that are used to clearly distinguish the two sets of high and low creatively played extracts in Sec 5.3.2., in order to elucidate areas of the brain associated with playing more or less creatively with sLORETA.

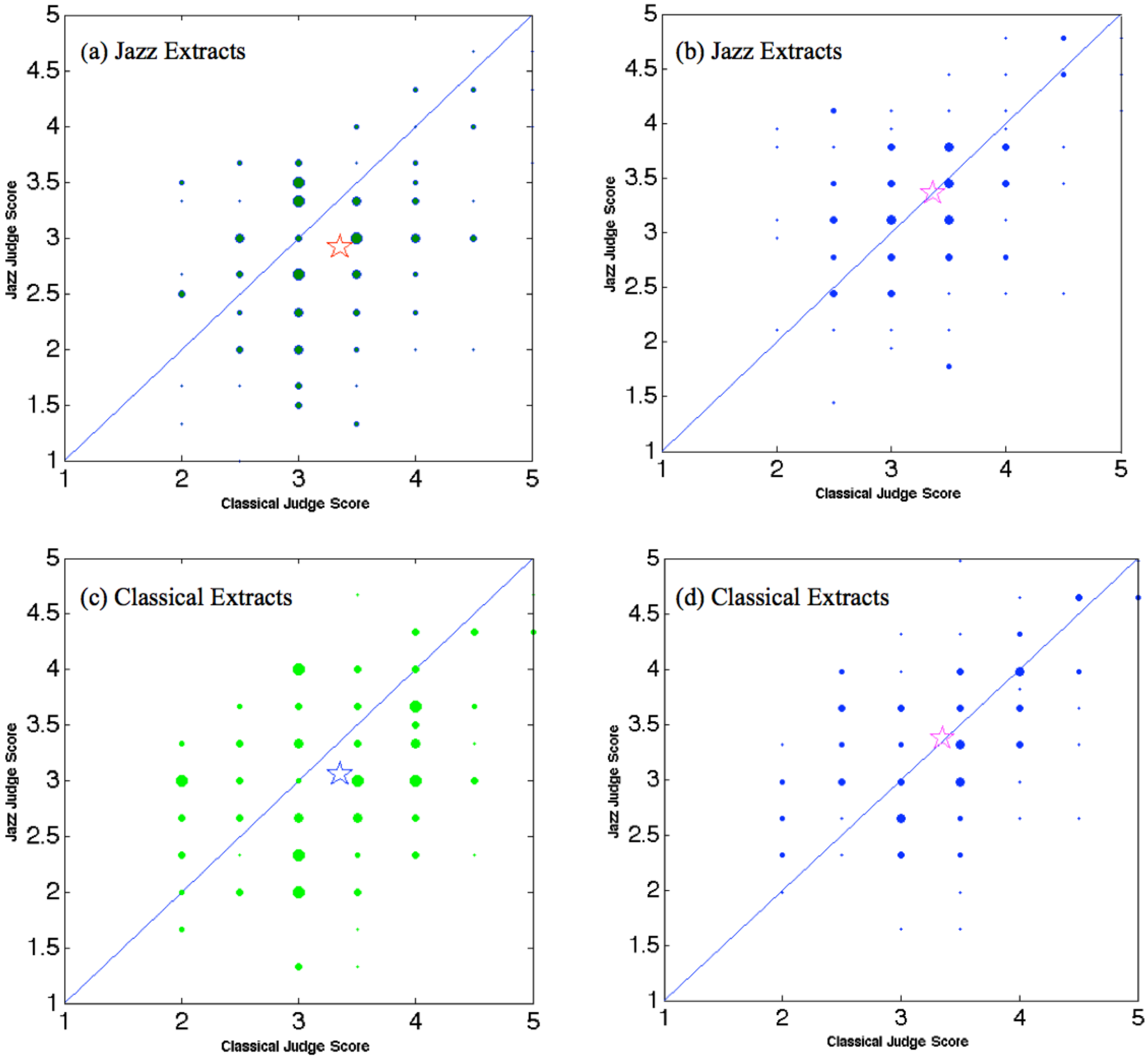


**Fig 1.2** (a) The scatter plot depicts mean scores given by Jazz judges versus Classical judges for Jazz extracts with the larger markers for more instances of that combination of judge scores. The centre of mass (shown by the star marker) indicates that Jazz judges give lower scores than Classical judges. (b) After correction, the scatter plot shows the centre of mass now lies on line with x=y with the rest of the combination scores better balanced on either side. (c) The scatter plot depicts mean scores given by Jazz judges versus Classical judges for Classical extracts again with the centre of mass showing that Jazz judges give lower scores than Classical judges. (d) The plot after correction with a balanced set of mean score combinations. (Because there are two Classical Judges, their mean values are in units of 1⁄2 while for the three Jazz Judges, their mean values are in units of 1/3).

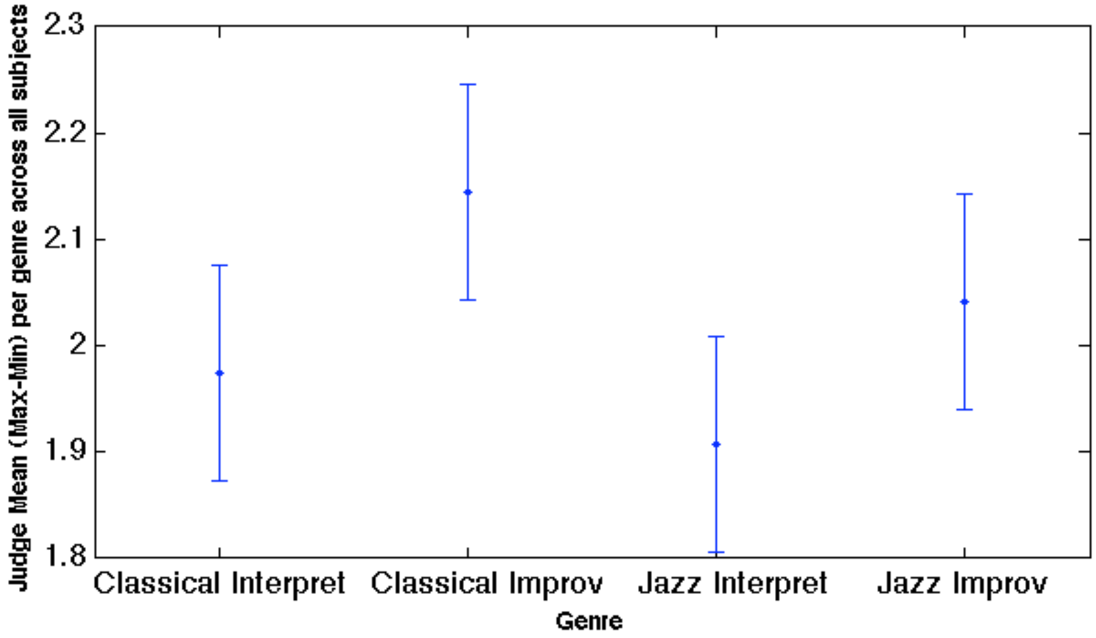


**Figure 1.3** The corrected mean of ‘Max-Min’ values with the overall mean of Judges assessments in red overhead, for the different genres and tasks of the experiment: Classical Interpretation, Classical Improvisation, Jazz Interpretation and Jazz Improvisation. We note that the Judges’ do not differ significantly in their agreement of assessment of extracts performed from the different genres or during the different tasks.

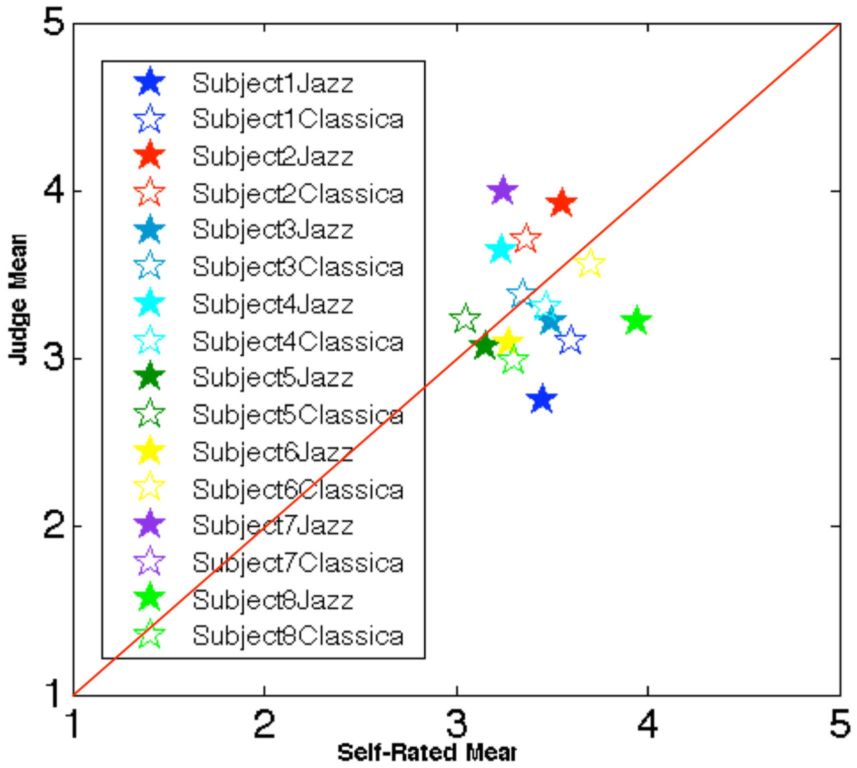


**Figure 1.4** Scatter plot of mean judge scores versus mean participants’ self-assessment separately across Jazz and Classical extracts.

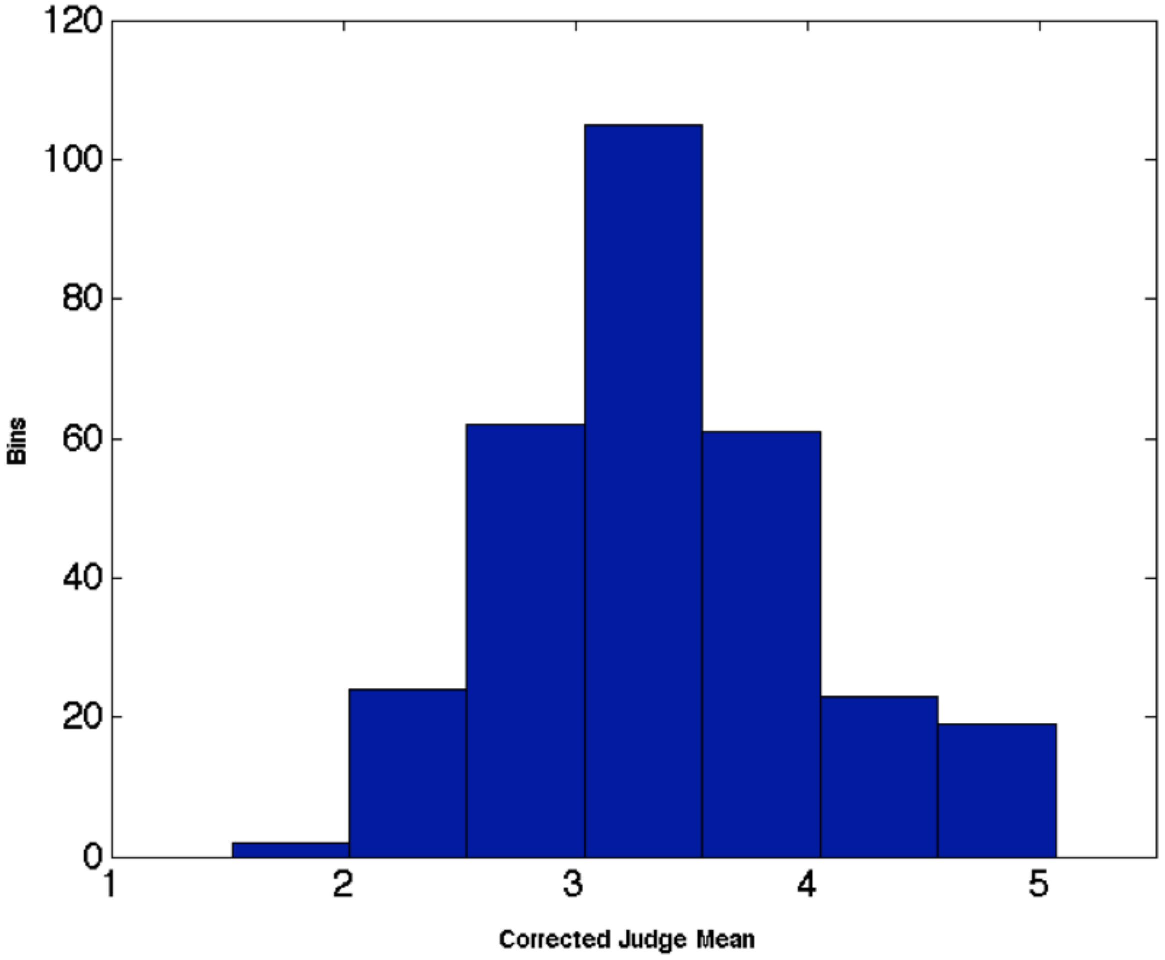


**Fig 1.5** Distribution of Judge Means (corrected) across all Subjects and Extracts (Normal Distribution) with random number generator

We performed standard multivariate analysis on the judges' and participants' assessments datasets using the Pearson's Correlation Coefficient, r:

$$r = \frac{\sum_{i=1}^{n}(X_i - \bar{X})(Y_i - \bar{Y})}{\sqrt{\sum_{i=1}^{n}(X_i - \bar{X})^2}\sqrt{\sum_{i=1}^{n}(Y_i - \bar{Y})^2}}$$

where X and Y are the values of two different assessment datasets e.g. X could be Judge 1 and Y could be Judge 2. We cross-correlated all 5 judges with each other and then subsequently each judge with each of the 8 participants across all extracts (correlation matrix C1), jazz extracts (correlation matrix C2) and classical extracts (correlation matrix C3).

| | 1 | 2 | 3 | 4 | 5 | 6 | 7 | 8 | 9 | 10 | 11 | 12 | 13 |
|---|---|---|---|---|---|---|---|---|---|---|---|---|---|
| 1 | 1 | 0.2598 | 0.2171 | 0.1387 | 0.2407 | 0.1279 | 0.4121 | −0.3694 | 0.1034 | 0.2495 | 0.0354 | 0.4100 | 0.0399 |
| 2 | 0.2598 | 1 | 0.5026 | 0.3183 | 0.1108 | 0.2150 | 0.5360 | −0.0637 | 0.4084 | −0.0451 | 0.1276 | 0.3685 | 0.4399 |
| 3 | 0.2171 | 0.5026 | 1 | 0.1870 | 0.4086 | 0.4294 | 0.4073 | −0.1570 | 0.1806 | 0.0723 | −0.1070 | 0.4323 | 0.3957 |
| 4 | 0.1387 | 0.3183 | 0.1870 | 1 | 0.2540 | −0.2119 | 0.2521 | −0.1812 | −0.4275 | 0.4038 | 0.0301 | 0.5587 | 0.0923 |
| 5 | 0.2407 | 0.1108 | 0.4086 | 0.2540 | 1 | 0.3176 | 0.3797 | 0.3925 | 0.3264 | 0.1698 | −0.1507 | 0.1501 | 0.5984 |
| 6 | 0.1279 | 0.2150 | 0.4294 | −0.2119 | 0.3176 | 1 | 0 | 0 | 0 | 0 | 0 | 0 | 0 |
| 7 | 0.4121 | 0.5360 | 0.4073 | 0.2521 | 0.3797 | 0 | 1 | 0 | 0 | 0 | 0 | 0 | 0 |
| 8 | −0.3694 | −0.0637 | −0.1570 | −0.1812 | 0.3925 | 0 | 0 | 1 | 0 | 0 | 0 | 0 | 0 |
| 9 | 0.1034 | 0.4084 | 0.1806 | −0.4275 | 0.3264 | 0 | 0 | 0 | 1 | 0 | 0 | 0 | 0 |
| 10 | 0.2495 | −0.0451 | 0.0723 | 0.4038 | 0.1698 | 0 | 0 | 0 | 0 | 1 | 0 | 0 | 0 |
| 11 | 0.0354 | 0.1276 | −0.1070 | 0.0301 | −0.1507 | 0 | 0 | 0 | 0 | 0 | 1 | 0 | 0 |
| 12 | 0.4100 | 0.3685 | 0.4323 | 0.5587 | 0.1501 | 0 | 0 | 0 | 0 | 0 | 0 | 1 | 0 |
| 13 | 0.0399 | 0.4399 | 0.3957 | 0.0923 | 0.5984 | 0 | 0 | 0 | 0 | 0 | 0 | 0 | 1 |

**Fig 1.6** Correlation matrix C1 cross-correlating judges and participants across all extracts. Columns/ rows 1 to 5 correspond to the 5 judges and columns/rows 6 to 13 correspond to the 8 participants.

| | 1 | 2 | 3 | 4 | 5 | 6 | 7 | 8 | 9 | 10 | 11 | 12 | 13 |
|---|---|---|---|---|---|---|---|---|---|---|---|---|---|
| 1 | 1 | 0.2598 | 0.2171 | 0.1387 | 0.2407 | −0.0363 | 0.0404 | −0.4804 | −0.0054 | 0.1967 | 0.1378 | 0.4217 | −0.3814 |
| 2 | 0.2598 | 1 | 0.5026 | 0.3183 | 0.1108 | −0.1206 | 0.3916 | −0.6727 | 0.4007 | 0.0077 | 0.2072 | 0.2296 | −0.4472 |
| 3 | 0.2171 | 0.5026 | 1 | 0.1870 | 0.4086 | 0.5028 | 0.0618 | −0.6405 | −0.0781 | −0.0662 | −0.0406 | 0.3358 | 0.1270 |
| 4 | 0.1387 | 0.3183 | 0.1870 | 1 | 0.2540 | −0.2607 | 0.4399 | −0.2325 | −0.4701 | 0.4031 | 0.0421 | 0.5798 | 0 |
| 5 | 0.2407 | 0.1108 | 0.4086 | 0.2540 | 1 | 0.0184 | 0.3306 | 0.3737 | 0.3705 | 0.0251 | −0.2941 | 0.2656 | 0.7127 |
| 6 | −0.0363 | −0.1206 | 0.5028 | −0.2607 | 0.0184 | 1 | 0 | 0 | 0 | 0 | 0 | 0 | 0 |
| 7 | 0.0404 | 0.3916 | 0.0618 | 0.4399 | 0.3306 | 0 | 1 | 0 | 0 | 0 | 0 | 0 | 0 |
| 8 | −0.4804 | −0.6727 | −0.6405 | −0.2325 | 0.3737 | 0 | 0 | 1 | 0 | 0 | 0 | 0 | 0 |
| 9 | −0.0054 | 0.4007 | −0.0781 | −0.4701 | 0.3705 | 0 | 0 | 0 | 1 | 0 | 0 | 0 | 0 |
| 10 | 0.1967 | 0.0077 | −0.0662 | 0.4031 | 0.0251 | 0 | 0 | 0 | 0 | 1 | 0 | 0 | 0 |
| 11 | 0.1378 | 0.2072 | −0.0406 | 0.0421 | −0.2941 | 0 | 0 | 0 | 0 | 0 | 1 | 0 | 0 |
| 12 | 0.4217 | 0.2296 | 0.3358 | 0.5798 | 0.2656 | 0 | 0 | 0 | 0 | 0 | 0 | 1 | 0 |
| 13 | −0.3814 | −0.4472 | 0.1270 | 0 | 0.7127 | 0 | 0 | 0 | 0 | 0 | 0 | 0 | 1 |

**Fig 1.7** Correlation matrix C2 cross-correlating judges and participants across jazz extracts. Columns/ rows 1 to 5 correspond to the 5 judges and columns/rows 6 to 13 correspond to the 8 participants.

| | 1 | 2 | 3 | 4 | 5 | 6 | 7 | 8 | 9 | 10 | 11 | 12 | 13 |
|---|---|---|---|---|---|---|---|---|---|---|---|---|---|
| 1 | 1 | 0.2598 | 0.2171 | 0.1387 | 0.2407 | 0.1791 | 0.6144 | −0.2877 | 0.3875 | 0.2928 | −0.1696 | 0.3737 | 0.3277 |
| 2 | 0.2598 | 1 | 0.5026 | 0.3183 | 0.1108 | 0.4496 | 0.6321 | 0.3879 | 0.4297 | −0.0300 | −0.0946 | 0.4192 | 0.6854 |
| 3 | 0.2171 | 0.5026 | 1 | 0.1870 | 0.4086 | 0.3728 | 0.5949 | 0.2133 | 0.5204 | 0.2457 | −0.2850 | 0.4174 | 0.4553 |
| 4 | 0.1387 | 0.3183 | 0.1870 | 1 | 0.2540 | −0.2017 | 0.0611 | −0.1200 | −0.4214 | 0.5021 | −0.0166 | 0.4784 | 0.1626 |
| 5 | 0.2407 | 0.1108 | 0.4086 | 0.2540 | 1 | 0.5309 | 0.4640 | 0.4464 | 0.3197 | 0.2929 | −0.1130 | 0.0469 | 0.5272 |
| 6 | 0.1791 | 0.4496 | 0.3728 | −0.2017 | 0.5309 | 1 | 0 | 0 | 0 | 0 | 0 | 0 | 0 |
| 7 | 0.6144 | 0.6321 | 0.5949 | 0.0611 | 0.4640 | 0 | 1 | 0 | 0 | 0 | 0 | 0 | 0 |
| 8 | −0.2877 | 0.3879 | 0.2133 | −0.1200 | 0.4464 | 0 | 0 | 1 | 0 | 0 | 0 | 0 | 0 |
| 9 | 0.3875 | 0.4297 | 0.5204 | −0.4214 | 0.3197 | 0 | 0 | 0 | 1 | 0 | 0 | 0 | 0 |
| 10 | 0.2928 | −0.0300 | 0.2457 | 0.5021 | 0.2929 | 0 | 0 | 0 | 0 | 1 | 0 | 0 | 0 |
| 11 | −0.1696 | −0.0946 | −0.2850 | −0.0166 | −0.1130 | 0 | 0 | 0 | 0 | 0 | 1 | 0 | 0 |
| 12 | 0.3737 | 0.4192 | 0.4174 | 0.4784 | 0.0469 | 0 | 0 | 0 | 0 | 0 | 0 | 1 | 0 |
| 13 | 0.3277 | 0.6854 | 0.4553 | 0.1626 | 0.5272 | 0 | 0 | 0 | 0 | 0 | 0 | 0 | 1 |

**Fig 1.8** Correlation matrix C3 cross-correlating judges and participants across jazz extracts. Columns/ rows 1 to 5 correspond to the 5 judges and columns/rows 6 to 13 correspond to the 8 participants.

Amongst the judges, the highest cross-correlation values were between judges 3 and 2, and 3 and 5, whilst the lowest cross-correlation values were between judges 2 and 5, and 1 and 4 (see figure 4). Amongst the judge/ participant cross-correlations, the highest positive correlations were between judge 5 and participant 8 across all and jazz extracts and between judge 2 and 8 for classical extracts. The highest anti-correlations were between judge 4 and participant 4 across all and classical extracts and between judge 2 and 3 for jazz extracts. Zero correlations (or near zero) were found between judge 4 and participant 6 for all and classical extracts and between judge 4 and participant 8 for jazz extracts (see figure 5 for judge/participant comparisons).

As can be seen, due to the low number of data points (down to 40 points for the judge/participant comparisons) and their discreteness this traditional statistical method may not be optimal for the main purposes of our study i.e. relating the assessments to the EEG recorded to ascertain high and non-creative extracts across all participants. Neither does it aid in quantitatively distinguishing accurately the participants' jazz or classical backgrounds whereas we used a simple method to deduce the phase space occupied by the combined judge and participant scores per genre for each participant, below or above their corresponding mid- line which makes them fall in one or the other genre category.

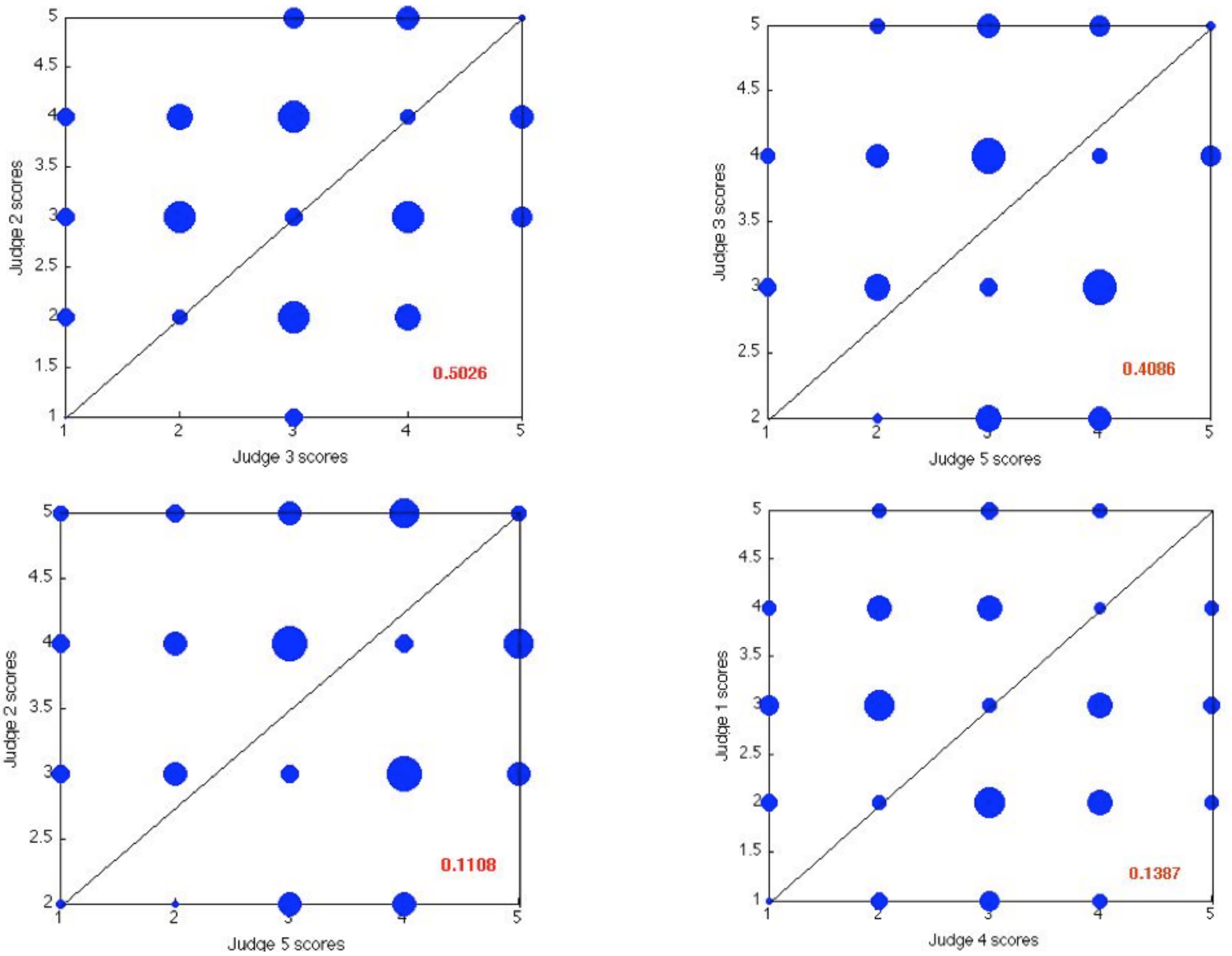


**Fig 1.9** Scatter plots of judge scores with their correlation coefficient in red. The top row consists of the two highest positive cross-correlations and the bottom row the two lowest.

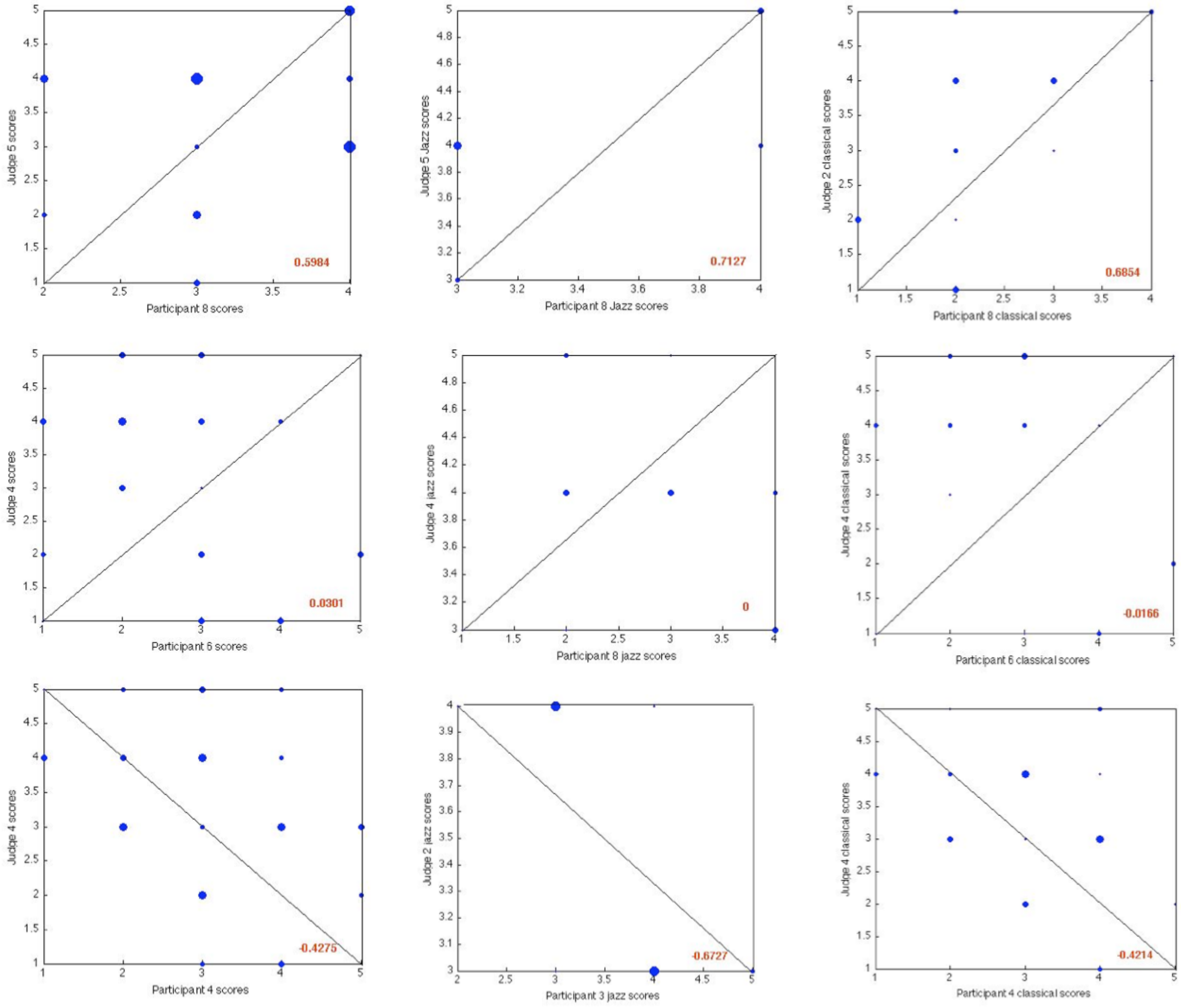


**Fig 1.10** Scatter plots of judge and participant scores with their correlation coefficient in red. The top row consists of the three highest positive cross-correlations, the middle row the three most non- correlated and the bottom row the three highest negative cross-correlations.

A Principal Component Analysis (PCA) was performed on each of the correlation matrices in order to orthogonally transform them into a set of values of linearly uncorrelated variables or ‘principal components’ that are equal to the original number of variables i.e. 13. In order to detect any patterns of clustering between subsets of judges and/or participants, we created a corresponding Euclidean distance matrix y for each correlation matrix from which a clustering matrix was calculated using a nearest neighbours method. These were then visualised using dendrograms (see fig 6). There seem to be fewer major clusters for judges across jazz extracts implying a uniformity in assessments further supported by figure 5.4 and the plots of (Max-Min) for the jazz genre across the interpretation and improvisation tasks.

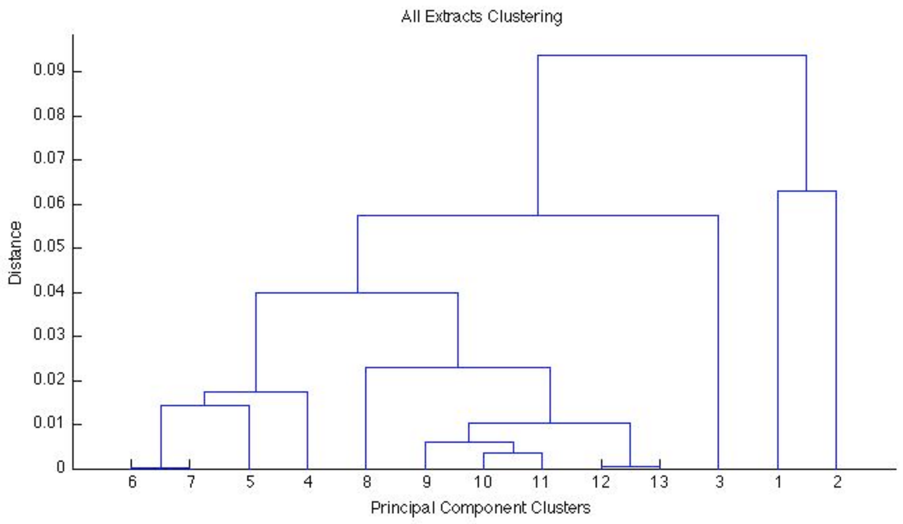


a)

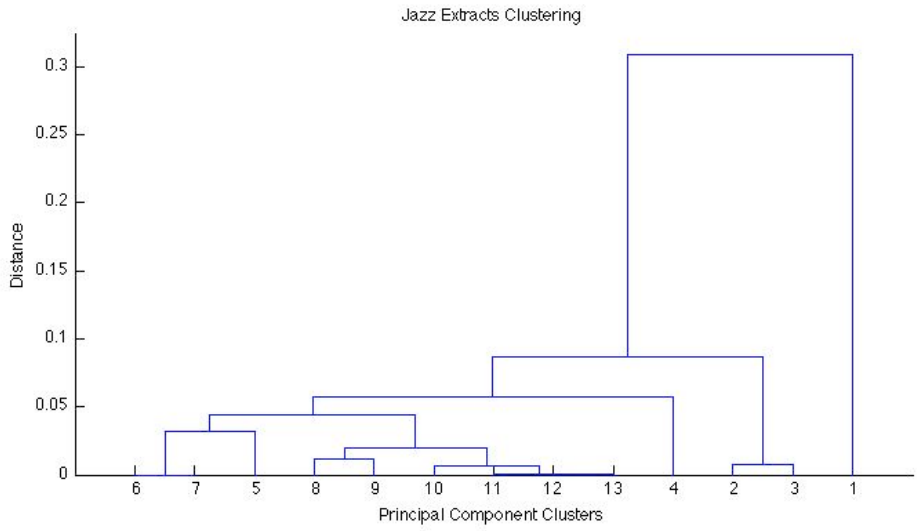


b)

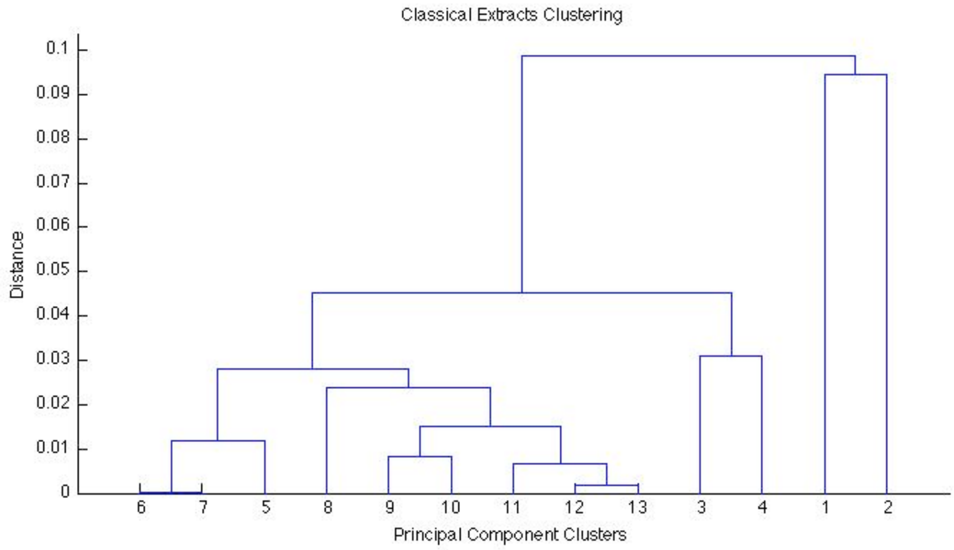


**Fig 1.11** Dendrograms of principal component clusters from correlation matrices for a) all extracts, b) jazz extracts and c) classical extracts. Components 1 to 5 correspond to 5*5 judge cross-correlations and components 6 to 13 correspond to 5*8 judge/participant cross-correlations.
